# Recurrent Composite Markers of Cell Types and States

**DOI:** 10.1101/2023.07.17.549344

**Authors:** Xubin Li, Justin Nguyen, Anil Korkut

## Abstract

Biological function is mediated by the hierarchical organization of cell types and states within tissue ecosystems. Identifying interpretable composite marker sets that both define and distinguish hierarchical cell identities is essential for decoding biological complexity, yet remains a major challenge. Here, we present RECOMBINE, an algorithm that identifies recurrent composite marker sets to define hierarchical cell identities. Validation using both simulated and biological datasets demonstrates that RECOMBINE achieves higher accuracy in identifying discriminative markers compared to existing approaches, including differential gene expression analysis. When applied to single-cell data and validated with spatial transcriptomics data from the mouse visual cortex, RECOMBINE identified key cell type markers and generated a robust gene panel for targeted spatial profiling. It also uncovered markers of CD8^+^ T cell states, including GZMK^+^HAVCR2^−^ effector memory cells associated with anti–PD-1 therapy response, and revealed a rare intestinal subpopulation with composite markers in mice. Finally, using data from the Tabula Sapiens project, RECOMBINE identified composite marker sets across a broad range of human tissues. Together, these results highlight RECOMBINE as a robust, data-driven framework for optimized marker selection, enabling the discovery and validation of hierarchical cell identities across diverse tissue contexts.

## Introduction

Tissue ecosystems comprise hierarchical cell types and dynamic states that underpin biological structure and function^1,2^. Characterizing these cell types and states has been a fundamental goal in biology. A key challenge lies in identifying a minimal set of composite markers capable of distinguishing hierarchical cell identities, particularly when these identities are defined by overlapping molecular features^3-5^. Such concise, composite markers, which capture the drivers of tissue heterogeneity, can facilitate the prediction of healthy and aberrant phenotypes, including immune states, developmental stages, and therapeutic responses.

Advances in single-cell transcriptomics have revealed the intricate cellular composition and organization of tissue ecosystems in both healthy and diseased contexts^6-10^. However, robust analytical approaches are needed to decode the composite markers that drive cellular heterogeneity and function within tissues. In parallel, the rise of targeted spatial transcriptomics technologies, including MERFISH^11^, STARmap^12^, and CosMx^13^, highlights the necessity of preselected marker panels to distinguish diverse cell types and states within tissues. Such marker panels are also critical for developing cost-effective clinical assays that accurately reflect biological processes underlying disease mechanisms^14^. Despite this pressing need, the identification of concise, composite markers capable of optimally discriminating hierarchical cell subpopulations remains a significant challenge.

Current computational approaches for identifying markers of cell subpopulations typically rely on partitional clustering to group cells into discrete clusters, followed by supervised marker selection to distinguish each cluster from others^15,16^. One widely used approach involves statistical tests to generate lists of differentially expressed genes (DEGs) between a specific cell cluster and other clusters. However, these DEG lists are often lengthy and challenging to interpret. Moreover, the selected markers are inherently biased by predefined cluster labels and fail to capture dynamic cell states within clusters. This limitation highlights a critical challenge in single-cell analysis: the ability to flexibly define varying levels of granularity for mapping cell identities^17^. Recently, partition-free methods have shown promise in addressing this challenge. For instance, MetaCell^18^ and SEACells^19^ create metacells that represent highly granular and distinct cell states, while Milo leverages K-nearest neighbor graph representations to identify perturbed cell states obscured by partitional clustering^20^. Despite these advances, a partition-free method capable of optimally selecting composite markers to characterize hierarchical cell identities remains an unmet need.

To address this challenge, we developed a novel computational framework, RECOMBINE (Recurrent Composite Markers for Biological Identities with Neighborhood Enrichment), designed to identify optimized, composite marker sets that distinguish hierarchical cell subpopulations. We assessed RECOMBINE’s performance using both simulated and biological datasets. Simulations demonstrated high precision and recall, while evaluations on biological data showed that RECOMBINE outperformed traditional differential gene expression (DEG) methods in characterizing hierarchical cell identities. Applied to single-cell and spatial transcriptomics data, RECOMBINE identified marker sets that: (i) delineated heterogeneous cell populations across cortical layers in the mouse visual cortex, (ii) characterized CD8^+^ T cell states, including one associated with immunotherapy response, (iii) distinguished a rare cell population in the mouse intestine, and (iv) captured the diversity of cell types across more than 20 human tissues. These results underscore the robustness and versatility of RECOMBINE in optimizing marker selection for hierarchical cell identities within complex tissue ecosystems.

## Results

### RECOMBINE identifies composite marker sets for hierarchical cell identities

RECOMBINE is a computational framework designed for unbiased selection of composite markers to distinguish hierarchical cell subpopulations (Figure 1A). It processes high-dimensional data, such as single-cell transcriptomics, to identify markers that delineate cell populations into hierarchical subgroups using the sparse hierarchical clustering with spike-and-slab lasso (SHC-SSL) algorithm. The SHC-SSL analysis produces two key outputs: (1) discriminant markers, identified as features with non-zero weights, and (2) cell similarities based on these discriminant markers. Building on these features, RECOMBINE identifies recurrent composite markers at the single-cell level through the construction of K-nearest neighbor graphs, followed by a neighborhood recurrence test. A neighborhood Z-score is calculated for each discriminant marker, reflecting the relative marker expression across all neighborhoods. Marker significance is determined using p-values derived from these Z-scores, with corrections for multiple hypothesis testing. To classify a marker as recurrent for a given cell subpopulation, two criteria are applied: the fraction of significant cells and the average neighborhood Z-score. The outputs of RECOMBINE include a panel of markers that distinguish cells, markers associated with the cell tree hierarchy, and recurrent composite markers specific to cell subpopulations.

**Figure 1.**
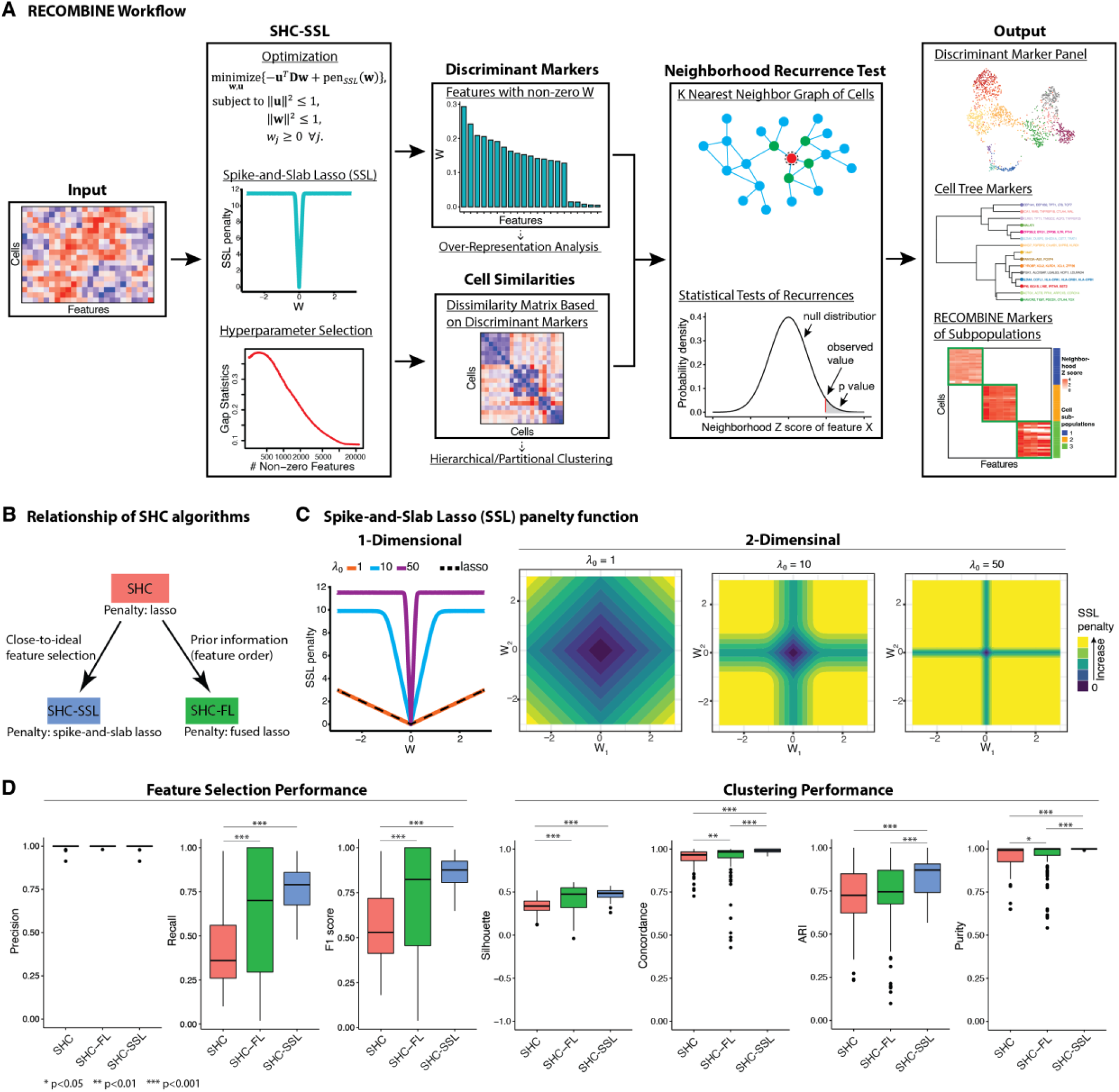
RECOMBINE: a computational framework that determines recurrent composite markers of cell types and states. **(A)** RECOMBINE workflow. **(B)** Relationship of sparse hierarchical clustering algorithms. **(C)** The SSL penalty function. The SSL penalty as a function of **w** with λ_1_ = 0.001 and λ_0_ ∈ {1,10,50}, where **w** is a one-dimensional (left panel) or two-dimensional variable (other panels). The SSL penalty converges to the lasso penalty with decreasing λ_0_ and to the L0-norm penalty with increasing λ_0_. Therefore, the SSL penalty forms a data-adjustable bridge between an L0-norm and a lasso penalty, which empowers SHC-SSL without any a priori information. **(D)** Benchmarking of the sparse hierarchical clustering algorithms for feature selection precision and recall, and clustering performance, using 100 randomly generated simulation datasets. Silhouette measures consistency of dissimilarity matrix based on discriminant features and ground truth labels of clusters, concordance measures clustering performance after hierarchical clustering, and adjusted Rand index (ARI) and purity measure clustering performance after Leiden clustering. For each metric, a higher value is better.

Using simulation data, we systematically benchmarked the performance of SHC-SSL against two other sparse hierarchical clustering algorithms: the conventional sparse hierarchical clustering with regular lasso (SHC)^21^ and a newly developed algorithm, SHC with fused lasso (SHC-FL) (Figure 1B). The SSL penalty in SHC-SSL provides a continuum between lasso and L0-norm, enabling more precise selection of discriminant features compared to the lasso used in SHC (Figure 1C). SHC-FL incorporates prior information about feature order through the fused lasso (Methods). All three algorithms achieved median precisions of 100%, indicating that the identified features were true discriminant markers (Figure 1D). SHC-SSL demonstrated the highest median recall and F1 score with minimal variation, reflecting superior feature selection accuracy and a low false-negative rate (Figure 1D). Incorporating prior information through SHC-FL significantly improved its feature selection performance. However, SHC-FL exhibited high recall variability (ranging from 20% to 100%), suggesting strong dependence on the alignment of prior information with the data. In contrast, SHC consistently showed lower recall and F1 scores compared to the other algorithms. To evaluate clustering performance, we assessed four metrics: silhouette score, concordance, adjusted Rand index, and purity (Methods). Across all metrics, performance scores increased monotonically from SHC to SHC-FL to SHC-SSL, with SHC-SSL exhibiting the best median performance and the least variation (Figure 1D). These findings demonstrate that SHC-SSL is the most accurate and robust algorithm for both feature selection and clustering among the tested sparse hierarchical clustering methods.

We next evaluated the performance of hierarchical clustering using RECOMBINE markers in comparison to DEGs (Figure S1). The analysis included three murine hematopoiesis datasets^66-68^ and one C. elegans embryogenesis dataset^69^. Cell labels were obtained from the original publications, except for the Dahlin et al. dataset, where cell annotations were unavailable, and unsupervised clustering based on all genes was used to generate cluster labels. For each dataset, RECOMBINE was employed to select an optimized set of discriminant markers from the complete gene set, while DEGs were identified through statistical tests between unsupervised clusters and subsequently filtered to match the total number of RECOMBINE markers. By comparing UMAP projections based on all genes, RECOMBINE markers, and DEGs, we observed that RECOMBINE markers effectively preserved the embeddings of the full gene set, whereas DEGs often failed to replicate similar embeddings (Figure S1). For instance, UMAP projections based on DEGs showed disconnected hematopoietic progenitor cells (Figure S1B) and compressed developmental trajectories (Figures S1C-D). To quantitatively assess the discriminative power of RECOMBINE markers relative to DEGs, we employed the gap statistic, a metric that measures the hierarchical clustering strength of selected markers relative to random markers of the same size. Across all tested datasets, RECOMBINE markers consistently exhibited stronger hierarchical clustering performance than DEGs (Figure S1). Collectively, our comprehensive evaluation using simulation and biological data demonstrates that RECOMBINE enables robust marker selection for hierarchical cell subpopulations.

### RECOBMINE optimizes marker panels for targeted spatial transcriptomics

Targeted spatial transcriptomics technologies have enabled spatially resolved molecular profiling at single-cell resolution^11-13^. However, these technologies require the prior selection of a gene panel capable of discriminating diverse cell types and states. To address this challenge, we utilized RECOMBINE, which selects markers from scRNA-seq data without any prior knowledge of cell type content or a pre-set number of markers. We applied RECOMBINE to an scRNA-seq dataset of the mouse visual cortex^26^ and validated the selected markers using a STARmap spatial transcriptomics dataset of the same tissue^12^. The scRNA-seq dataset included 14,249 cells representing 23 cell types across three major compartments: excitatory neurons, inhibitory neurons, and non-neuronal cells. RECOMBINE identified 366 discriminant genes (Figure 2A, S2A, Table S1), including key markers for inhibitory neurons (e.g., Gad1, Gad2, Vip, Npy, Synpr, Cck, Sst, and Rab3b) and excitatory neurons (e.g., Slc17a7, Pcp4, and Nrgn). Additionally, RECOMBINE identified recurrent composite markers that delineate the cellular hierarchy of each cell type (Figure 2B, S2C). The UMAP projection of cells based on RECOMBINE markers successfully separated all 23 cell types (Figure 2C). Hierarchical clustering strength based on RECOMBINE markers was significantly higher compared to that based on DEGs (Figure 2D). While all RECOMBINE markers were included in the DEG set, the number of RECOMBINE markers (N=366) represented only 12% of the total DEGs (N=3,096), demonstrating optimized discriminative power with substantially fewer markers (Figure 2E). Notably, RECOMBINE markers were enriched in highly significant DEGs but also included markers from less significant DEGs, suggesting that these less significant genes may play crucial roles in distinguishing hierarchical subpopulations (Figure S2B).

**Figure 2.**
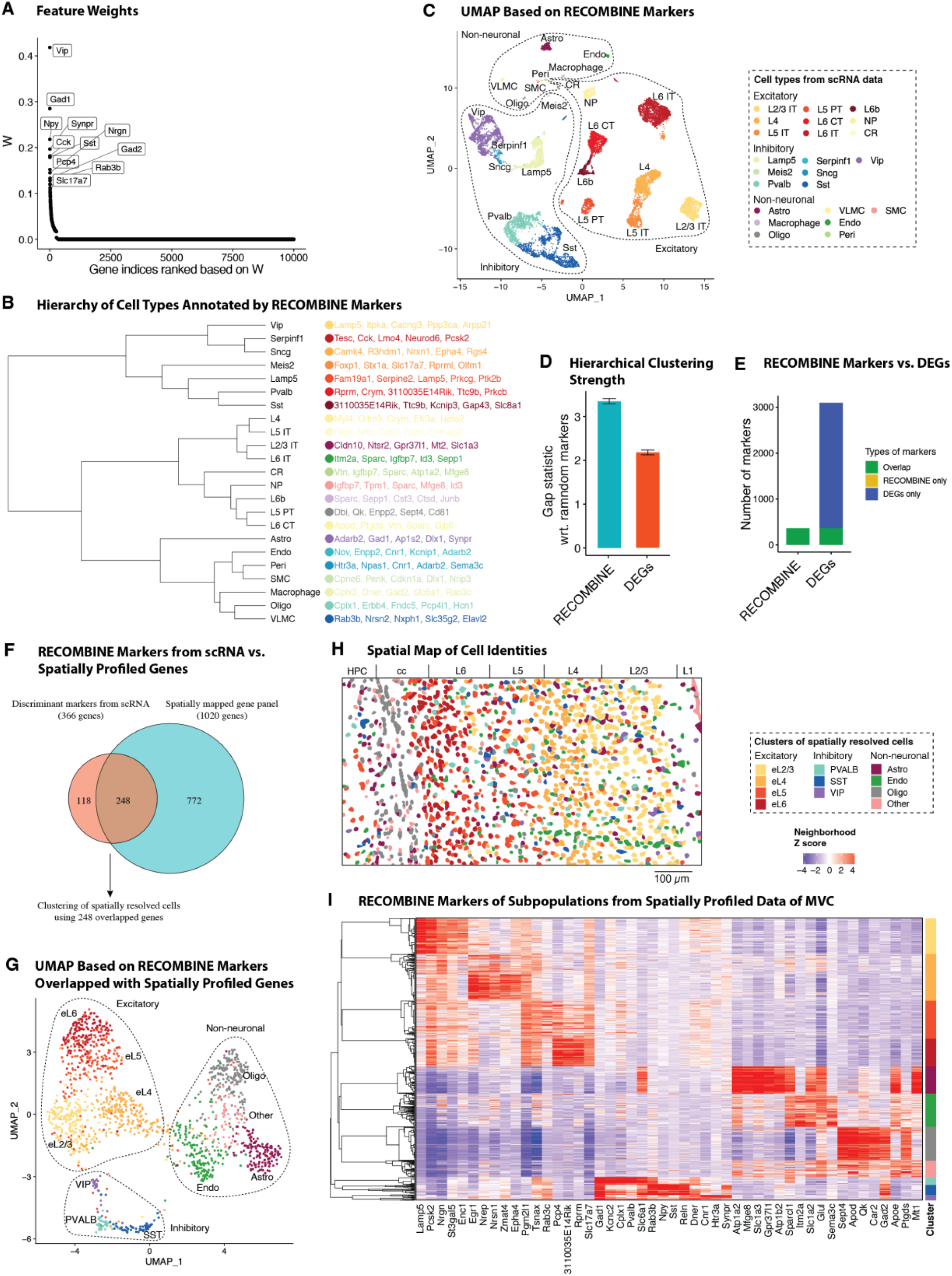
RECOMBINE selects composite markers that define hierarchical cell types and provides an optimized panel for spatial profiling of mouse visual cortex. **(A)** Feature weights of all genes in which the top 10 discriminant markers are labeled. **(B)** Hierarchy of cell types annotated by top 5 RECOMBINE markers. **(C)** UMAP of cells based on RECOMBINE markers, where colors indicate cell type labels from Tasic et al. (Nature, 2018). Cell type abbreviations: IT, intratelencephalic; PT, pyramidal tract; CT, corticothalamic; NP, near-projecting; CR, Cajal– Retzius cell; Astro, astrocyte; Oligo, oligodendrocyte; VLMC, vascular leptomeningeal cell; Endo, endothelial cell; Peri, pericyte; and SMC, smooth muscle cell. **(D)** Comparison of RECOMBINE markers and DEGs. DEGs were obtained by Wilcoxon tests of each cell type with respect to the rest of cell types and filtered based on absolute log_2_ fold-change > 0.25 and false discovery rate < 0.05. **(E)** Comparison of hierarchical clustering performance between RECOMBINE markers and top DEGs of the same size. **(F)** Venn diagram of discriminant markers in the scRNA data and the genes profiled in the spatial data. **(G)** Heatmap of neighborhood Z scores showing gene modules across cell clusters of the spatial data. **(H)** UMAP of spatially resolved cells based on RECOMBINE markers (N=248) and colored by clusters. **(I)** Spatial map of cells with the same color code of cell clusters in (E). L1-6, the six neocortical layers; cc, corpus callosum; HPC, hippocampus.

The hierarchical tissue structure and associated marker sets within the mouse cortex were decoded with a bottom to top analysis on the RECOMBINE output (Figure 2B, Figure S2D). Starting from the bottom branches on the cell population tree, we computed the RECOMBINE features that are shared by the cells contained at each branch of the tree (Figure 2B). Next, we moved up to a parental node to compute the shared features for the corresponding cell population. This was continued until reaching to a cell hierarchy where cell populations diverged and did not carry any shared RECOMBINE features. This approach enabled us to define hierarchical cell populations with lower hierarchies connected through as well as disconnected hierarchies at higher levels with no shared feature. The application to the cortex tissue identified three distinct cell type groups matched to excitatory, inhibitory, and non-neuronal compartments. Within each group, the connected and hierarchical cell identities shared common markers (Figure S2D). While the cell types in each cluster were hierarchically linked to each other through the shared markers (Figure S2D), they were disconnected from the cell types in the other groups.

A comparison between the 366 RECOMBINE markers and 1,020 genes profiled in the STARmap dataset revealed 248 overlapping markers (Figure 2F). Using these overlapped markers, unsupervised clustering of STARmap data identified 11 cell clusters across the excitatory neuron, inhibitory neuron, and non-neuronal compartments (Figure 2G). Spatial mapping of these clusters recapitulated the layered organization of excitatory neurons and the sparse distribution of inhibitory neurons (Figure 2H). The RECOMBINE markers formed distinct modules across cell clusters and defined cell subpopulations, while some markers were shared across clusters or compartments, reflecting the continuous nature of gene expression across cell types (Figure 2I, S2E). For example, Slc6a1 and Gad2 were enriched in inhibitory neurons but also expressed in astrocytes and oligodendrocytes, respectively. We hypothesized that the most enriched markers could form a reduced-size panel with minimal loss of discriminative power. To test this, we selected the top 10 enriched markers from each cluster, resulting in a panel of 69 markers (Figure 2I). UMAP projections using these 69 markers effectively separated all cell subpopulations, closely mirroring the patterns observed with the full set of discriminant markers, except for cluster eL4 being farther from cluster eL5 (Figure S2F). These results demonstrate that RECOMBINE enables unbiased, data-driven selection of marker panels from paired scRNA-seq data, providing a robust foundation for targeted spatial transcriptomics.

### RECOMBINE defines CD8 T cell states and markers predictive of cancer immunotherapy response

Advancing cancer immunotherapies, such as immune checkpoint blockade and adoptive cell therapy, requires a comprehensive understanding of CD8 T cell states and their associated markers^27-30^. Using RECOMBINE, we identified composite markers of CD8 T cell states from a pan-cancer scRNA-seq dataset comprising 234,550 CD8 T cells^31^. RECOMBINE selected 320 discriminant markers that effectively distinguished subgroups of CD8 T cells (Figure 3A, S3A, and Table S2). Among the top 10 ranked markers were cytotoxic molecules (e.g., FGFBP2, FCGR3A, NKG7, GZMA, GZMB, and GZMH) and cytokine- and growth factor-related molecules (e.g., CCL5, CCR6, CCR7, and FGFBP2), reflecting effector and memory functions. Hierarchical clustering strength based on RECOMBINE markers was higher than that based on DEGs (Figure S3B). Unsupervised clustering of RECOMBINE markers revealed 15 clusters (Figure 3B, S3C), each characterized by composite markers that were either unique to specific clusters or shared among clusters with similar CD8 T cell states (Figure 3B–C). Examination of checkpoint gene expression highlighted cluster c10, which exhibited elevated levels of both stimulatory (e.g., CD27, TNFRSF9, TNFRSF18) and inhibitory (e.g., CTLA4, PDCD1, LAG3, HAVCR2, TIGIT) checkpoint molecules, suggesting a terminally exhausted state (Figure 3D). Furthermore, cluster c10 showed enriched expression of KIR2DL4, an inhibitory checkpoint molecule typically expressed on NK cells, aligning with recent findings that an NK cell–like transition is a hallmark of CAR T cell dysfunction^32^. In summary, RECOMBINE accurately identified CD8 T cell states and their characteristic markers, providing valuable insights for improving cancer immunotherapies.

**Figure 3.**
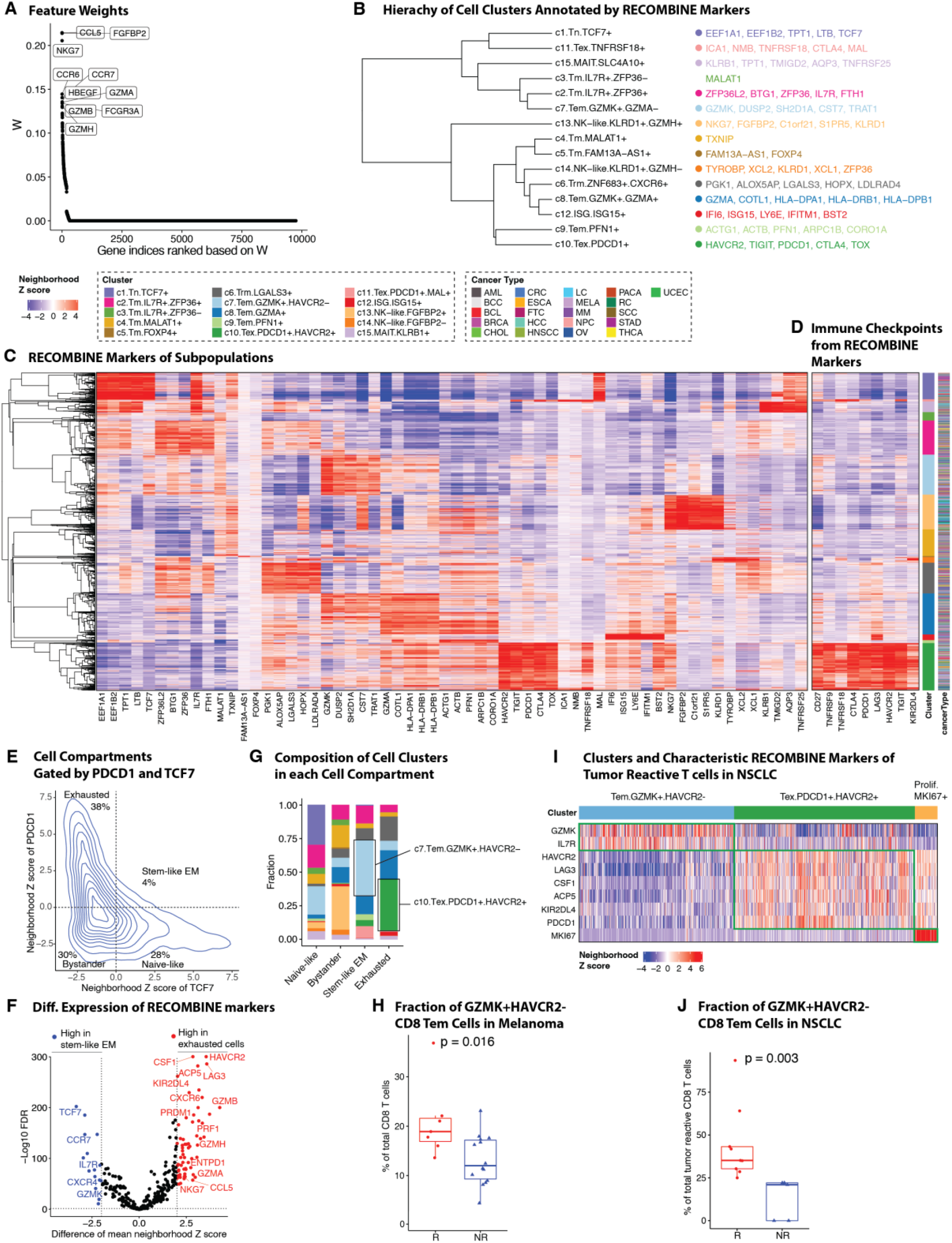
RECOMBINE identifies CD8 T cell states and markers associated with cancer immunotherapy response. **(A)** Feature weights of all genes in which the top 10 discriminant markers are labeled. **(B)** Hierarchy of cell clusters annotated by top 5 RECOMBINE markers. **(C)** Heatmap of neighborhood Z scores showing gene modules across clusters. **(D)** Heatmap of neighborhood Z scores showing stimulatory and inhibitory immune checkpoint genes identified by RECOMBINE. **(E)** CD8 T cell compartments gated by neighborhood Z scores of TCF7 and PDCD1. EM: effector memory. **(F)** Differentially expressed markers between stem-like EM and exhausted T cell compartments defined in (E). **(G)** Compositions of cell clusters across the cell compartments defined in (E). **(H)** Comparison of GZMK^+^HAVCR2^-^ T_em_ (c7 in RECOMBINE analysis) CD8 T cell fractions in immunotherapy responders (R, N=7) vs. non-responders (NR, N=14) of melanoma patients treated with anti–PD-1 therapy (Sade-Feldman et al. 2018). **(I)** Heatmap of neighborhood Z scores showing characteristic markers of clusters of 13,403 tumor-reactive CD8 T cells from anti-PD-1–treated and treatment-naïve patients with non-small cell lung cancer (Liu et al. 2022). **(J)** Comparison of GZMK^+^HAVCR2^-^ T_em_ cell fractions within tumor-reactive CD8 T cells in anti–PD-1 therapy responder (R, N=9) vs. non-responder (NR, N=5) patients with non-small cell lung cancer.

We next investigated whether characteristic markers and CD8 T cell subpopulations were associated with responses to immunotherapies, such as PD-1 blockade. Previous studies have indicated that exhausted CD8 T cells (TCF7^−^PDCD1^+^) acquire a stable and distinct epigenetic profile that remains minimally remodeled after anti–PD-1 therapy^33,34^. In contrast, stem-like effector memory (EM) CD8 T cells (TCF7^+^PDCD1^+^) provide tumor control and exhibit a proliferative burst in response to anti–PD-1 therapy^35-37^. To identify characteristic markers and specific cell subpopulations responsive to anti–PD-1 therapy, we classified all CD8 T cells into four compartments—naïve-like, bystander, stem-like EM, and exhausted cells—based on neighborhood Z-scores of TCF7 (a stemness marker) and PDCD1 (an activation and exhaustion marker) (Figure 3E). To distinguish stem-like EM cells from exhausted cells, we conducted differential analyses of the neighborhood Z-scores for discriminant markers between these two populations (Figure 3F, Table S2). As anticipated, stem-like EM cells were significantly enriched in naïve and memory markers (e.g., TCF7, CCR7, IL7R, CXCR4), whereas exhausted cells were enriched in cytotoxic and terminal differentiation markers (e.g., GZMA, GZMB, GZMH, PRF1, NKG7, CCL5, CXCR6, ENTPD1, PRDM1). Interestingly, GZMK expression was significantly elevated in stem-like EM cells, suggesting a role in early-stage EM cell development, while inhibitory checkpoint molecules HAVCR2, LAG3, and KIR2DL4 (but not PDCD1, CTLA4, or TIGIT) were highly enriched in exhausted cells, implicating these molecules in the late-stage development of terminally exhausted cells. In alignment with these findings, cluster c7 (GZMK^+^HAVCR2^−^, representing Tem cells) and cluster c10 (PDCD1^+^HAVCR2^+^, representing Tex cells) were identified as the largest subpopulations of stem-like EM and exhausted cells, respectively (Figure 3G). These results highlight distinct markers and subpopulations that may serve as potential targets for improving anti–PD-1 immunotherapy efficacy.

As stem-like EM CD8 T cells are likely the primary targets of anti–PD-1 therapy, we hypothesized that GZMK^+^HAVCR2^−^ Tem cells might play a pivotal role in mediating therapeutic responses. To test this, we analyzed the association of GZMK^+^HAVCR2^−^ Tem cells with clinical outcomes in a subset of anti–PD-1-treated melanoma patients (N=21) described by Sade-Feldman et al. ^38^ Consistent with our hypothesis, the enrichment of c7 (GZMK^+^HAVCR2^−^ Tem) cells was significantly higher in patients who responded to anti–PD-1 therapy (Figure 3H). To validate this finding, we applied RECOMBINE to an independent scRNA-seq dataset comprising 13,403 tumor-reactive CD8 T cells from both anti–PD-1-treated and treatment-naïve patients with non-small cell lung cancer (NSCLC; N=34)^39^. Three major cell states were identified with characteristic markers: GZMK^+^HAVCR2^−^ Tem, PDCD1^+^HAVCR2^+^ Tex, and MKI67^+^ proliferative cells (Figure 3I). Among post-therapy samples from this cohort (N=14), the GZMK^+^HAVCR2^−^ Tem cell population was significantly enriched in responders, consistent with the findings in melanoma patients (Figure 3J). Furthermore, the top markers of exhausted cells (e.g., HAVCR2, LAG3, KIR2DL4, CSF1, and ACP5) identified in the pan-cancer dataset (Figure 3F) were recapitulated in the NSCLC dataset (Figure 3I), reinforcing their roles in T cell exhaustion. Notably, KIR2DL4, CSF1, and ACP5 remain less characterized in the context of T cell exhaustion, warranting further investigation to uncover their potential contributions. In conclusion, RECOMBINE analyses identified characteristic markers of stem-like EM and exhausted CD8 T cells, demonstrated the enrichment of GZMK^+^HAVCR2^−^ Tem cells in responders to anti–PD-1 therapy, and proposed novel markers of T cell exhaustion. These findings provide a foundation for future studies aimed at improving cancer immunotherapy by targeting CD8 T cell states and their associated markers.

### RECOMBINE discovers composite markers for rare cell subpopulations

To highlight RECOMBINE’s ability to identify rare cell types, a critical goal in single-cell omics analysis, we analyzed an scRNA-seq dataset of mouse intestinal organoids^40^. Hierarchical clustering based on 183 RECOMBINE markers identified five distinct clusters: enterocytes, two clusters of enterocyte precursor cells, transit-amplifying cells, and secretory cells (Figure 4A–B, S4A, and Table S3). The top 10 markers effectively discriminated the common enterocyte subpopulation from other cell subpopulations. Within the enterocyte precursor cluster, two subpopulations were identified: late-stage cells, which were more differentiated toward enterocytes, and early-stage cells, which were closer to transit-amplifying cells. RECOMBINE further identified three gene modules corresponding to enterocytes, transit-amplifying cells, and secretory cells (Figure 4C). Over-representation analysis of RECOMBINE markers revealed that transit-amplifying cells were enriched with genes encoding ribosomal proteins and mRNAs, whereas enterocytes were enriched with genes associated with metabolic pathways (hypergeometric test, FDR < 0.001) (Figure 4D). Notably, secretory cells within cluster c5 included a rare subpopulation of Reg4^+^ enteroendocrine cells, previously identified in Grün et al.^40^ This rare population was characterized by high expression of Reg4 and additional genes, including Cd24a, Actg1, Hopx, Jun, Tff3, Chgb, Egr1, Tac1, Chga, and Fos (Figure 4C, E). While Reg4, Chga, and Chgb have been used previously to characterize Reg4^+^ enteroendocrine cells^40^, these genes were also expressed in a fraction of enterocytes and enterocyte precursor cells, limiting their specificity as markers for this rare subpopulation (Figure 4C). To more selectively identify and fully characterize Reg4^+^ enteroendocrine cells, additional markers such as Tff3 or Tac1, as detected by RECOMBINE, should be included (Figure 4C).

**Figure 4.**
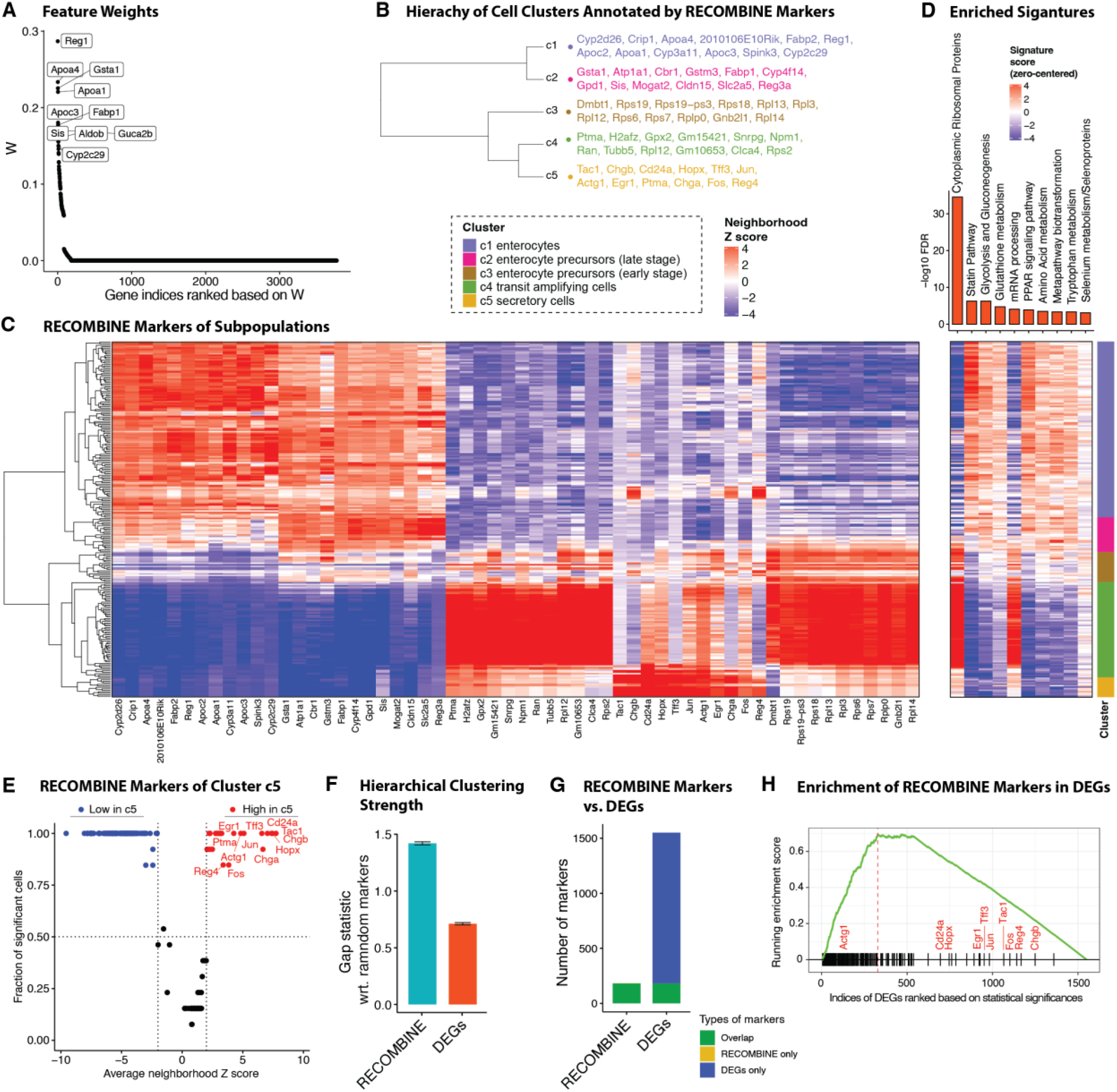
RECOMBINE identifies concise yet discriminative markers of a rare cell subpopulation of mouse intestine. **(A)** Feature weights of all genes where the top 10 discriminant markers are labeled. **(B)** Hierarchy of clusters annotated by top 10 RECOMBINE markers. **(C)** Heatmap of neighborhood Z scores showing gene modules across clusters. **(D)** Enrichment of mouse pathway signatures in RECOMBINE markers (top) and heatmap of signature scores across cells (bottom). Top 10 significantly enriched signatures are shown. **(E)** Volcano plot of cluster c5-specific markers, highlighting markers of a rare cell subpopulation in red. **(F)** Comparison of hierarchical clustering performance between RECOMBINE markers and top DEGs of the same size. **(G)** Comparison of RECOMBINE markers and DEGs. DEGs were obtained by Leiden clustering of all cells based on all genes followed by Wilcoxon test for each cluster with respect to the rest of the clusters. RECOMBINE markers were statistically significant markers with false discovery rate (FDR) < 0.05, and DEGs were filtered based on absolute log_2_ fold-change > 0.25 and FDR < 0.05. **(H)** Enrichment plot of RECOMBINE markers in the list of DEGs ranked by their significance from Wilcoxon tests.

We compared RECOMBINE markers to DEGs and observed that RECOMBINE markers exhibited greater hierarchical clustering strength (Figure 4F). While all but one RECOMBINE marker was included within the DEG set, the number of RECOMBINE markers (N=183) was significantly smaller than the total DEGs (N=1,550) (Figure 4G). Notably, RECOMBINE markers were enriched among the more significant DEGs, yet some, including nine of the 10 characteristic markers of Reg4^+^ enteroendocrine cells, were statistically less significant DEGs (Figure 4H, S4B). For instance, although Reg4 ranked lower in DEG significance, it exhibited much greater statistical significance in the RECOMBINE analysis. These findings demonstrate that RECOMBINE selects a concise yet highly sensitive set of markers, enabling the identification of rare Reg4^+^ enteroendocrine cells. This population required not only previously reported markers but also additional RECOMBINE-identified markers to achieve effective discrimination from other cell subpopulations.

### RECOMBINE identifies compact marker sets for diverse cell types across human tissues

Finally, we applied RECOMBINE to the Tabula Sapiens dataset, a human single-cell transcriptomic reference atlas comprising nearly 500,000 cells from over 20 tissues and organs^70^. This analysis aimed to evaluate RECOMBINE’s capacity to identify discriminant markers across heterogeneous cell contexts in a large-scale dataset while generating a comprehensive resource of tissue marker sets. Single-cell transcriptomics data and cell ontology class annotations were collected for tissues from the Tabula Sapiens (Figure 5A). For each tissue, RECOMBINE identified recurrent composite marker genes for cell ontology classes (Figure 5B and Table S4). To enable integrated analysis, we averaged expression data to generate pseudobulk expression matrices, which were then pooled across all tissues. UMAP projections based on RECOMBINE markers revealed that transcriptomic variation was primarily driven by cell lineages (Figure 5C) rather than tissue of origin (Figure 5D). Hierarchical clustering further demonstrated RECOMBINE’s ability to identify selective markers unique to specific cell types and shared functional markers spanning multiple lineages (Figure 5E). For example, well-known markers such as CD79A, MS4A1, and BANK1 were identified for B cells, while immunoglobulin genes (IGHG3, IGLC3, IGKC) were linked to plasma cells. Skeletal muscle cells exhibited selective expression of genes like ENO3, MYBPC1, CA3, MYOZ1, and ACTA1, whereas eye-specific cells, such as photoreceptors and Müller cells, uniquely expressed ROM1, AIPL1, SAG, CRX, and CNGB1. Additionally, shared markers included genes related to antigen presentation (HLA-DQA1, HLA-DPA1, CD74) found in B cells, macrophages, and dendritic cells, as well as stress-related heat shock protein genes (HSP90AB1, HSPE1, HSPA1A) spanning multiple lineages. In conclusion, RECOMBINE effectively identified both specific and shared markers across heterogeneous cell contexts, underscoring its utility for comprehensive human pan-tissue transcriptomic analyses.

**Figure 5.**
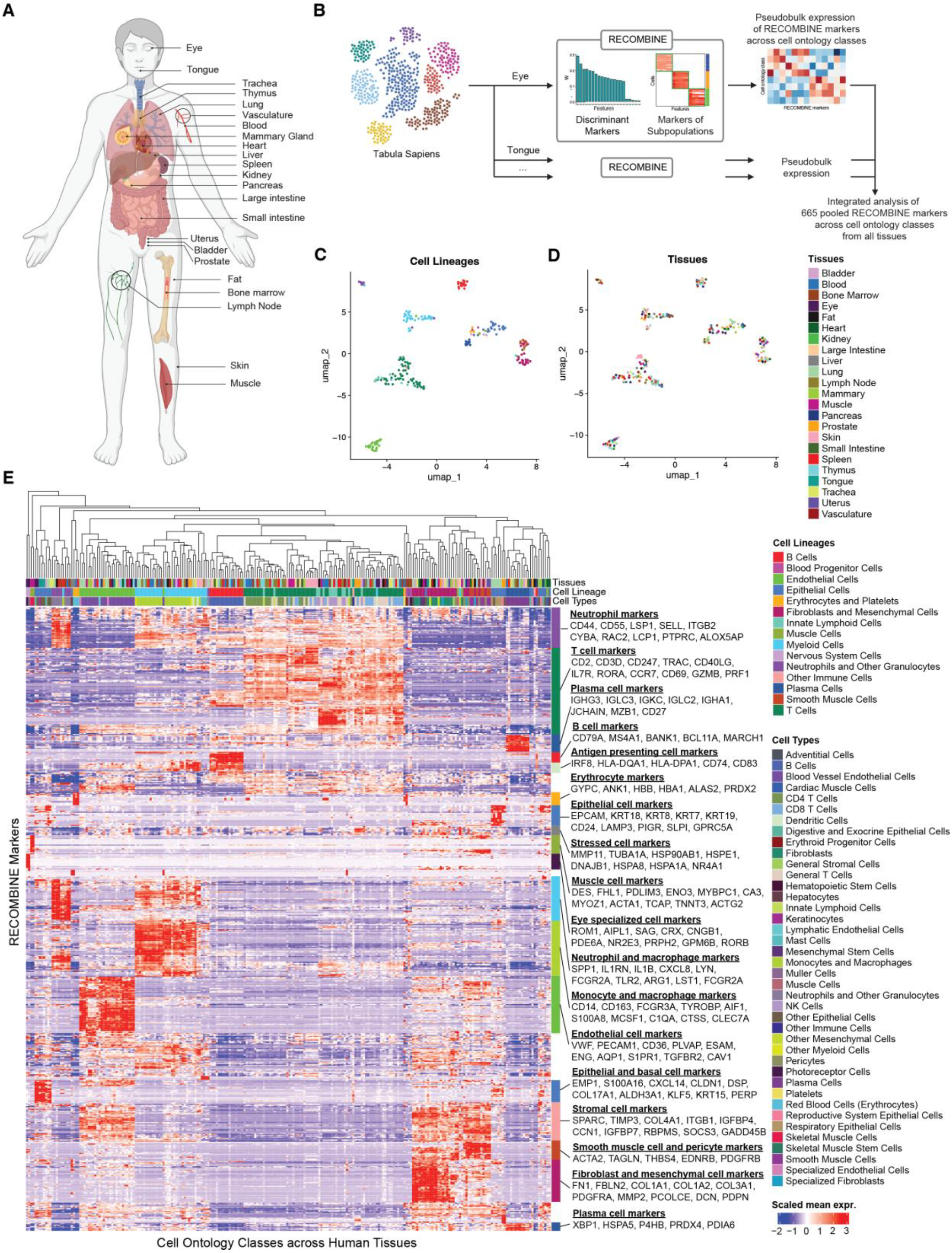
RECOMBINE identifies concise discriminant markers of cell types across human tissues. **(A)** Schematic of the tissues profiled by single-cell RNA sequencing from the Tabula Sapiens dataset used in this study. **(B)** Workflow of RECOMBINE applied to the Tabula Sapiens to identify recurrent composite markers that effectively discriminate cell types in all tissues. **(C-D)** UMAP projections of pseudobulk expression of RECOMBINE markers across cell types in all tissues, colored by cell lineage (C) and tissue type (D). **(E)** Heatmap showing pseudobulk expression of RECOMBINE markers across cell types in all tissues.

## Discussion

Here, we present the development and applications of RECOMBINE, a computational framework designed to select optimized and composite markers that discriminate hierarchical cell identities. RECOMBINE is built on the sparse hierarchical clustering with spike-and-slab lasso (SHC-SSL) algorithm, which outperformed the conventional sparse hierarchical clustering with lasso (SHC) in both optimal marker selection and accurate clustering, as demonstrated in our benchmarking studies. Evaluations across diverse biological datasets revealed that RECOMBINE markers exhibited superior cell population discrimination strength compared to differentially expressed genes (DEGs), a conventional approach for identifying cell subpopulations from single cell omics data. Unlike partition-based methods such as DEGs—which identify markers separating predefined clusters without accounting for the underlying structure of tissue and cell hierarchies—RECOMBINE couples marker selection with hierarchical clustering. This coupling allows RECOMBINE to identify composite markers that discriminate both cell types and dynamic states with high granularity. The simultaneous optimization of marker selection and hierarchical clustering enables RECOMBINE to detect markers across all levels of the cell and tissue hierarchy, including distinct cell types, continuous cell states, and rare cell subpopulations. Despite its simplicity, this unique capability underscores its utility for advancing our understanding of cellular heterogeneity in complex biological systems.

We demonstrated RECOMBINE’s versatility through its application across diverse biological domains. Using scRNA-seq data from the mouse visual cortex, RECOMBINE identified discriminant markers for cell types and states. These markers successfully delineated spatially resolved cell subpopulations in the same tissue using a STARmap dataset, highlighting RECOMBINE’s robust performance in marker selection for targeted spatial transcriptomics. In scRNA-seq data from pan-cancer CD8 T cells, RECOMBINE identified composite markers for CD8 T cell states, including GZMK^+^HAVCR2^−^ effector memory cells associated with anti–PD-1 therapy response, and novel markers related to CD8 T cell exhaustion. Similarly, in mouse intestinal tissue scRNA-seq data, RECOMBINE identified selective markers characterizing a rare subpopulation of Reg4^+^ enteroendocrine cells. Finally, RECOMBINE identified composite marker sets for heterogeneous cell types across more than 20 human tissues in the Tabula Sapiens. Taken together, RECOMBINE demonstrates broad applicability for data-driven extraction of recurrent composite marker sets to characterize diverse cell subpopulations across biological systems.

RECOMBINE introduces several key algorithmic advances over existing methods. First, RECOMBINE does not require a pre-specified marker panel size; instead, it determines an optimal size through a statistically principled and data-driven approach using the gap statistic. In contrast, methods like SCMER^46^ require the user to define the marker panel size a priori, which is often unknown and may lead to suboptimal selection for discriminating cell subpopulations. This adaptive approach ensures that RECOMBINE selects marker panels tailored to the underlying data. Second, RECOMBINE enforces marker sparsity using a spike-and-slab lasso penalty function, which has demonstrated superior marker selection performance compared to the lasso penalty function employed in methods like SHC, scGeneFit^47^, and RANKCORR^48^. Third, RECOMBINE incorporates neighborhood enrichment tests to identify marker modules associated with cell subpopulations, enabling biologically meaningful interpretations of cell types and states. Finally, a notable advantage of RECOMBINE is its ability to select an optimized and compact set of discriminant markers. Compared to DEGs, RECOMBINE produces a substantially smaller marker set while maintaining high sensitivity. This compactness facilitates easier interpretation and practical utility of the marker list but may result in some loss of comprehensiveness. As observed in our benchmark study, SHC-SSL achieved a median precision of 100% but a median recall of 79%. This imperfect recall can be attributed to the fact that some markers may be omitted if the selected markers already provide maximum discrimination for cell subpopulations.

RECOMBINE is a scalable algorithm capable of analyzing hundreds of thousands of cells, as demonstrated in the pan-cancer CD8 T cell analysis with 234,550 cells. Achieving this scalability required addressing the computational challenge inherent to RECOMBINE’s marker selection algorithm, SHC-SSL, which operates on a dissimilarity matrix of dimension *n*^2^ by *p*, (*n*: cells, *p*: features), leading to quadratic increases in memory usage as *n* grows. To overcome this limitation, we implemented a pseudo-cell strategy that merges cells with shared transcriptomic profiles into pseudo-cells and identifies markers that discriminate between these pseudo-cells. Provided the pseudo-cells represent fine-grained cell groups, this approach approximates the solution for selecting discriminant markers at the single-cell level. An additional benefit of the pseudo-cell strategy is its ability to reduce technical noise and stabilize variance, thereby enabling more robust marker selection. Notably, similar strategies have been employed by methods such as MetaCell and SEACells, which construct granular and distinct cell group representations to address sparsity in single-cell data while preserving fine-grained heterogeneity of cell states. The outputs from MetaCell and SEACells—representing granular cell groups—can also be seamlessly integrated into RECOMBINE as an alternative to the pseudo-cell strategy, serving as inputs for marker selection. With these strategies, along with ongoing advancements, we anticipate that RECOMBINE will remain highly scalable for applications in large-scale projects^49^.

RECOMBINE is a generalized framework for the unbiased selection of recurrent composite markers to characterize hierarchical biological identities. Its applications, as demonstrated through diverse examples, can uncover how multicellular systems are organized into hierarchical cell identities that mediate biological functions. By generating marker modules for cell subpopulations, RECOMBINE can identify drivers of key biological processes. For instance, it can detect immune cell state-specific marker sets for subpopulations implicated in resistance to immunotherapy. Moreover, RECOMBINE markers can inform the development of new assays for identifying unique processes, cell types, and dynamic states within tissues using single-cell omics data. While our studies focused primarily on single-cell expression profiles, RECOMBINE is adaptable to a variety of omics data modalities, including protein expression, epigenomic profiles, and genomic alterations, in both single-cell and bulk resolutions. This versatility positions RECOMBINE as a valuable tool for diverse biological analyses. Through its capacity to discover novel drivers of biological functions, characterize rare cell subpopulations in complex tissues, and guide strategies for manipulating biological processes (e.g., selection of therapeutic interventions), RECOMBINE has the potential to profoundly advance our understanding of biological systems and enhance our ability to manipulate their functions.

## Methods

### Overview of RECOMBINE

Given single-cell omics data, RECOMBINE uncovers recurrent composite marker sets of hierarchical cell identities in two steps. In the first step, RECOMBINE employs the SHC-SSL algorithm to select the markers that best discriminate cells while performing hierarchical clustering. The discriminant markers consist of a minimal set that characterize the data and can be subjected to over-representation analysis for biological interpretation. Besides hierarchical clustering, the cell dissimilarity matrix based on the discriminant markers can be used for hierarchical or partitional clustering. In the second step, RECOMBINE uses neighborhood recurrence tests to extract markers of cell subpopulations from the discriminant markers. Specifically, the cell dissimilarity matrix is used to build a K nearest neighbor graph of cells. For any individual cell, its nearest neighbors compose its local neighborhood context. Then, a neighborhood Z score is calculated to quantify the recurrence of each of the discriminant markers for each cell’s local neighborhood. Finally, based on statistical significances of neighborhood Z scores, markers are identified at the single cell level; based on fractions of cells having significant neighborhood Z scores, markers are identified at the subpopulation level.

In below sections, we first describe SHC-based algorithms by reviewing the previously developed SHC algorithm and introducing SHC-SSL and another related algorithm that we developed, SHC-FL. As these algorithms involve hyperparameters, we next describe a method for hyperparameter selection. Then, we give details of simulation data and metrics for benchmarking the three algorithms. Next, as SHC-SSL is computationally expensive when the sample size is large, we describe a pseudo-cell strategy for reducing its computational cost. Then, we introduce a neighborhood recurrence test for extracting cell-level and subpopulation-level markers from discriminant markers. Finally, we give details of applying RECOMBINE to diverse biological datasets.

Here, we describe some notations. For any vector **v** ∈ ℝ^*n*^, ‖**v**‖ denotes the L2 norm (Euclidian norm) of **v**, and ‖**v**‖_1_ denotes the L1 norm of **v**.

### Sparse hierarchical clustering

First, we summarize the previously developed SHC algorithm^21^, which we applied to identify recurrent oncogenic co-alterations^50^. Given an *n* × *p* data matrix **X** with *n* samples by *p* features, we denote *x*_*i*_ as a sample that is a *p*-dimensional vector indexed by *i*, where _1_ ≤ *i* ≤ *n* and *d*(*x*_*i*_, *x*_*i*_′) as a measure of the dissimilarity between samples *x*_*i*_ and *x*_*i*_′. Assuming the dissimilarity measure is additive in features, that is, *d*(*x*_*i*_, *x*_*i*_′) = ∑_*j*_ *d*_*i,i*_′_,*j*_, where *d*_*i,i*_′_,*j*_ is the dissimilarity measure between samples *x*_*i*_ and *x*_*i*_′ along feature *j*, the dissimilarity matrix becomes the solution of a constrained optimization problem:

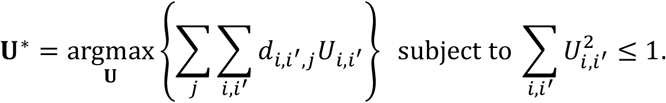

It can be shown as the optimal solution 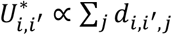. Thus, with this reformulation, hierarchical clustering performed on **U**^*^ is equivalent to that performed on {*d*(*x*_*i*_, *x*_*i*_′)}_*n*×*n*_. To calculate *d*_*i,i*_′_,*j*_, one could use the squared Euclidean distance *d*_*i,i*_′_,*j*_ = (*X*_*ij*_ − *X*_*i*_′_*j*_)^2^ or the absolute difference *d*_*i,i*_′_,*j*_ = |*X*_*ij*_ − *X*_*i*_′_*j*_|.

The weighted dissimilarity matrix was introduced in the SHC method. SHC provides a means to select a small number of discriminant features that contribute to a weighted dissimilarity matrix. Specifically, SHC solves the optimization problem

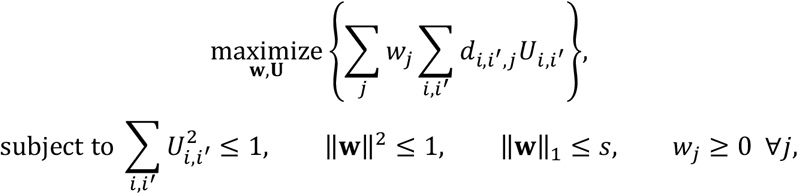

where **w** is a weight vector for features and *s* is a tuning parameter that controls sparsity of **w**. When *s* is small, the lasso, or L1, penalty of **w** encourages many elements of **w** to be zero. The L2 penalty of **w** is necessary because without it, only one element of **w** would be nonzero. Let **U**^**^ optimize the above criterion, from which we have 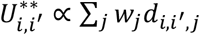, where the dissimilarity along each feature is weighted according to *w*_*j*_. Due to these constraints, a small value of *s* makes *w* sparse, so **U**^**^ depends only on a small subset of features. Therefore, clustering based on **U**^**^ depends only on the selected features.

To simplify representation, let’s rearrange {*d*_*i,i*_′_,*j*_} _*n*×*n*×*p*_ to get a matrix 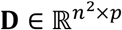 in which column *j* consists of ordered elements {*d*_*i,i*_′_,*j*_}_*n*×*n*_ stacked into a vector. Accordingly, let 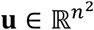 be the vector by stacking {*U*_*i j*_ ′} _*n*×*n*_. Then, the SHC optimization criterion is in the form

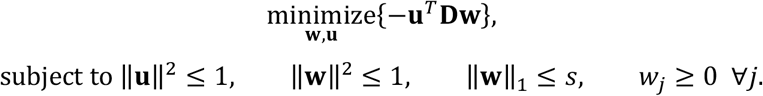

We can iteratively optimize **u**and **w**. With **w** fixed, this criterion is equivalent to

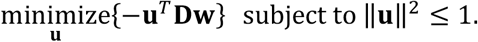

The optimal 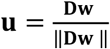, and *u*_*i*_ ≥ 0 for _1_ ≤ *i* ≤ *n*^2^ since all the elements of **D** and **w** are nonnegative. With **u** fixed, the criterion takes the form

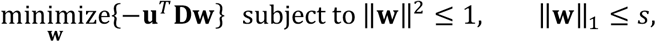

where we drop the non-negativity constraint because it is redundant when all the elements of **u** and **D** are non-negative. The optimal 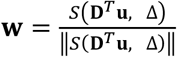, where *S* is the soft-thresholding operator *S*(*x, c*) = sign(*x*)(|*x*| − *c*)_+_, and Δ = 0 if this results in ‖**w**‖_1_ ≤ *s*; otherwise, Δ > 0 is chosen such that ‖**w**‖_1_ = *s*.

### Sparse hierarchical clustering with spike-and-slab lasso

Here, we introduce the SHC-SSL algorithm, which uses an SSL penalty. We start with a spike-and-slab Laplace prior distribution of **w**, from which an SSL penalty function follows^51,52^. Suppose the prior distribution of **w** is in the form

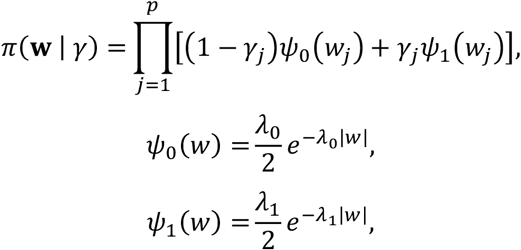

where **γ :**= (γ_*j*_) _1≤*j*≤*n*_, γ_*j*_ ∈ {0, 1}. Each *w*_*j*_ is a mixture of two Laplace distributions, *ψ*_0_ and *ψ*_1_, and γ_*j*_ is the latent variable indicating the mixture components to which *w*_*j*_ belongs. With λ_0_ large, *ψ*_0_(*w*) serves a restrictive, point-mass-like “spike distribution” for modeling a small *w*; in contrast, with λ_1_ small, *ψ*_1_(*w*) serves a diffusive, heavy-tailed “slab distribution” for modeling a large *w*. Suppose the latent variable **γ** has a hierarchical prior distribution as

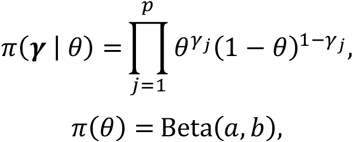

where *θ* is shared across γ_*j*_’s and Beta(*a, b*) is the beta distribution with shape parameters *a, b* > 0. As we can see, π(γ_*j*_ = _1_| *θ*) = *θ*; thus, *θ* acts as the prior probability of *w*_*j*_ being large. Since *θ* is shared across *w*_*j*_’s, intuitively *θ* reflects the expected fraction of large *w*_*j*_’s. By marginalizing out **γ**, we obtain

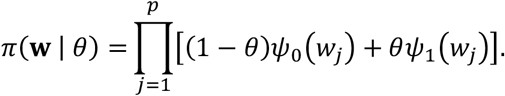

Furthermore, by marginalizing out *θ*, we have

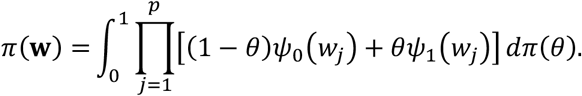

With this prior marginal distribution of **w**, we define the SSL penalty as

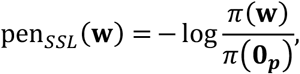

which is centered so that pen_*SSL*_ (**0**) = 0. Note that our definition of the SSL penalty is the negative of the one described by Ročková and George^51^. We choose this definition because it is consistent with the definition of the lasso penalty that is the negative logarithm of the Laplace distribution. When λ_0_ → ∞ and λ_1_ → 0, the SSL penalty is equivalent to an L0-norm penalty; when λ_0_ = λ_1_, it is equivalent to a lasso penalty. Therefore, the SSL penalty forms a continuous bridge between an L0-norm and a lasso penalty. We keep λ_1_ to a small constant and tune λ_0_ during hyperparameter selection.

Now, we replace the lasso with the SSL penalty in the objective of SHC, yielding the optimization criterion as

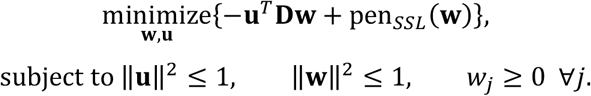

We iteratively optimize **u** and **w**. With **w** fixed, the optimal **u** is calculated in the same way as the one in SHC. With **u** fixed, the SHC-SSL objective takes the form

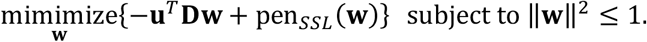

To find stationary points of the above objective, we rewrite it as a Lagrangian function

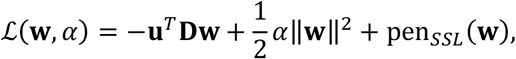

where α ≥ 0 is the Lagrange multiplier and a factor of 1/2 is included for later convenience. Following the sub-differential calculus, we obtain

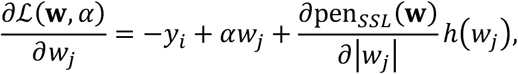

where **y** = **D**^*T*^**u** and *h*(*w*) is a subgradient of |*w*|, meaning *h*(*w*) = sign(*w*) when *w* ≠ 0, and *h*(*w*) ∈ [−1−1]when *w* = 0.

Let’s present the derivative of the SSL penalty. Using Lemma 2 and 3 described by Ročková and George^51^, the derivative of the SSL penalty with respect to |*w*_*j*_| is in the form

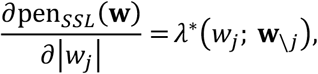

where

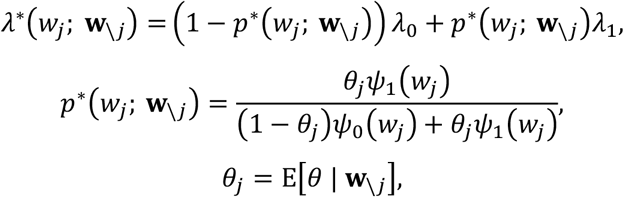

where E[*θ* | **w**_\*j*_]is the posterior expectation of *θ* given **w**_\*j*_. Here, λ^*^(*w*_*j*_; **w**_\*j*_) can be interpreted as a weighted average of λ_0_ and λ_1_, and the weight *p*^*^(*w*_*j*_; **w**_\*j*_) is the conditional probability of *w*_*j*_ being from *ψ*_1_(*w*_*j*_) rather than from *ψ*_0_(*w*_*j*_). Thus, a larger *w*_*j*_ tends to have a smaller λ^*^(*w*_*j*_; **w**_\*j*_). Both λ^*^(*w*_*j*_; **w**_\*j*_) and *p*^*^(*w*_*j*_; **w**_\*j*_) depend on *θ*_*j*_. To calculate *θ*_*j*_ = E[*θ* | **w**_\*j*_], note that when *p* is large, E[*θ* | **w**_\*j*_]should be similar to E[*θ* | **w**]. From Lemma 4 in Ročková and George^51^, *θ*_*j*_ can be approximated by

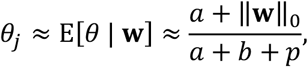

where ‖**w**‖_0_ is the L0 norm of **w**.

Now, we solve for the optimal **w**. Substituting the derivative of pen_*SSL*_ (**w**), the derivative of *L*(**w**, α) can be expressed as

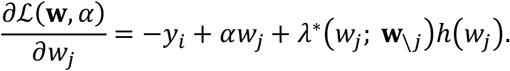

Let 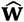 denote an estimate of the global optimum of **w**. At stationary points, with the Karush-Kuhn-Tucker conditions, 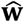 needs to satisfy

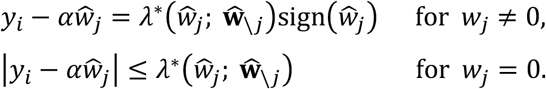

This can be written equivalently as

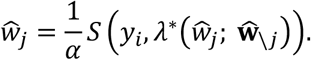

where the soft thresholding operator *S*(*x, c*) = sign(*x*)(|*x*| − *c*)_+_. At optimum, α is chosen such that 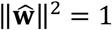.

It should be noted that the above necessary conditions are not sufficient for the global mode under the SSL penalty^51^. Using Theorem 3.1 from Ročková^52^, the global mode 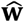 under the SSL penalty can be expressed as

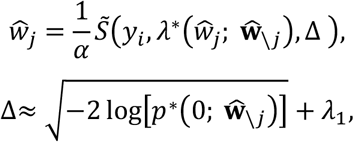

where 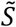 is a generalized thresholding operator defined as 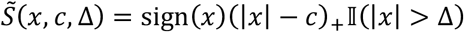 and α is chosen such that 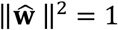. We can think of 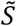 as a composition operator consisting of an L0-type hard thresholding operator *S*_0_(*x*, Δ) = *x*II(|*x*| > Δ) followed by a lasso-type soft thresholding operator *S*_1_(*x, c*) = sign(*x*)(|*x*| − *c*)_+_. Both operators adaptively depend on the value of 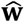. Each coordinate of 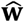 has its own self-adaptive shrinkage, which borrows information across other coordinates. The hard thresholding operator *S*_0_ depends on Δ, which in turn depends on 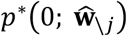, the conditional probability of 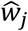 being zero is from the slab distribution. A larger 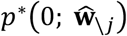, corresponding to a smaller Δ, induces a smaller hard thresholding effect on 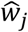 (i.e., *S*_0_ tends to leave 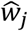 as is). In contrast, a larger hard thresholding effect is induced if 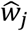 being zero is likely from the spike distribution (i.e., *S*_0_ tends to set 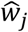 as 0). On the other hand, the soft thresholding operator *S*_1_ adaptively depends on 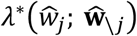. As we discussed before, a larger 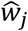 has a smaller 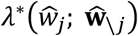. When 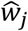 is large, it is shrunk by a small amount (close to λ_1_); when 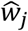 is small, it is shrunk by a large amount (close to λ_0_). Therefore, in contrast to the constant shrinkage of the lasso penalty, the SSL penalty induces a self-adaptive shrinkage that depends on itself and borrows information from other coordinates. In other words, while lasso constantly biases 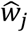 toward zero^53^, SSL shrinks strongly small 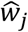 ‘s toward zeros but only introduces very slight biases for large 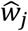 ‘s. For example, as shown in Fig. 1c, by fixing λ_1_ = 0.00_1_ and increasing λ_0_ from 1 to 50, the SSL penalty forms a continuum between a lasso and an L0-norm penalty. When λ_0_ = _1_, the SSL penalty function is hardly distinguishable from lasso. When λ_0_ = 50, the SSL penalty function has a sharp transition from zero to non-zero weights of features, approaching the L0-norm penalty that is known to be ideal for feature selection.

#### SHC-SSL algorithm

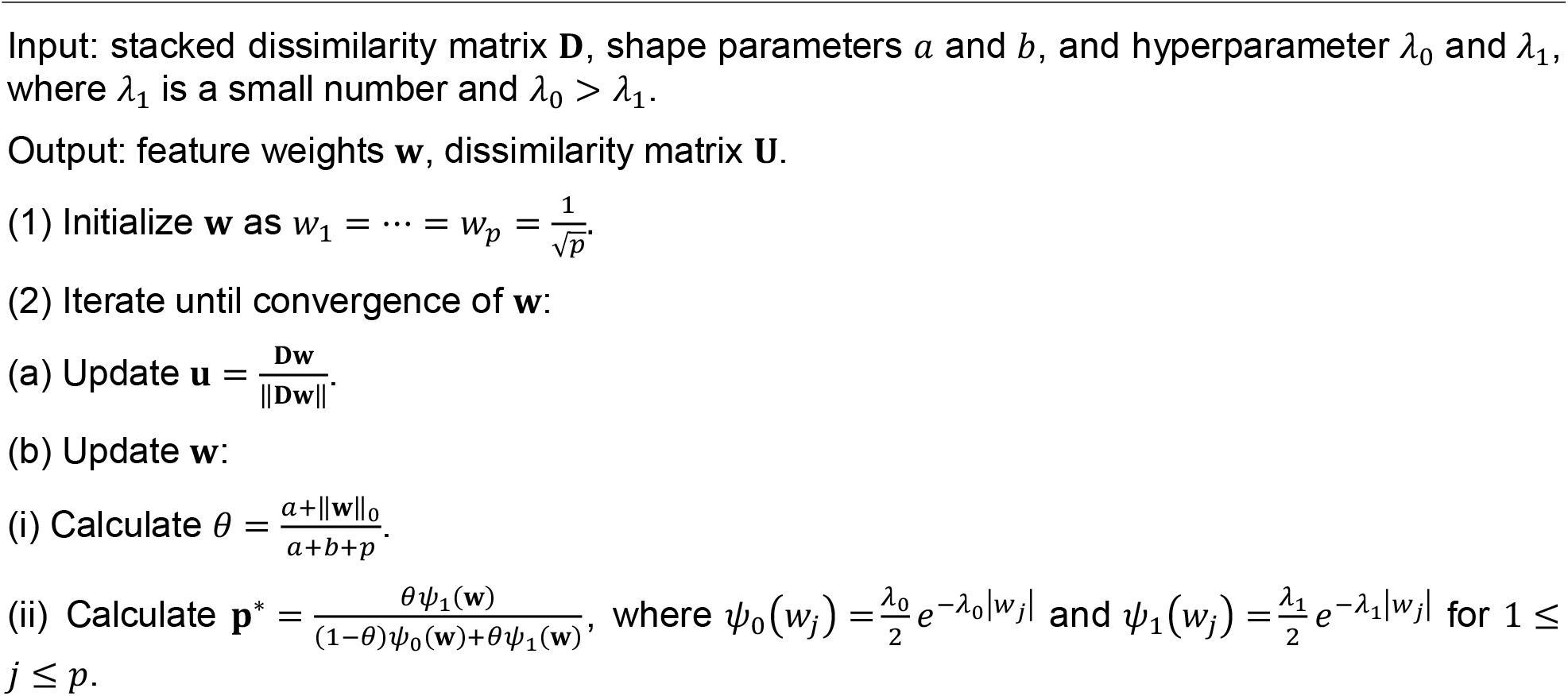
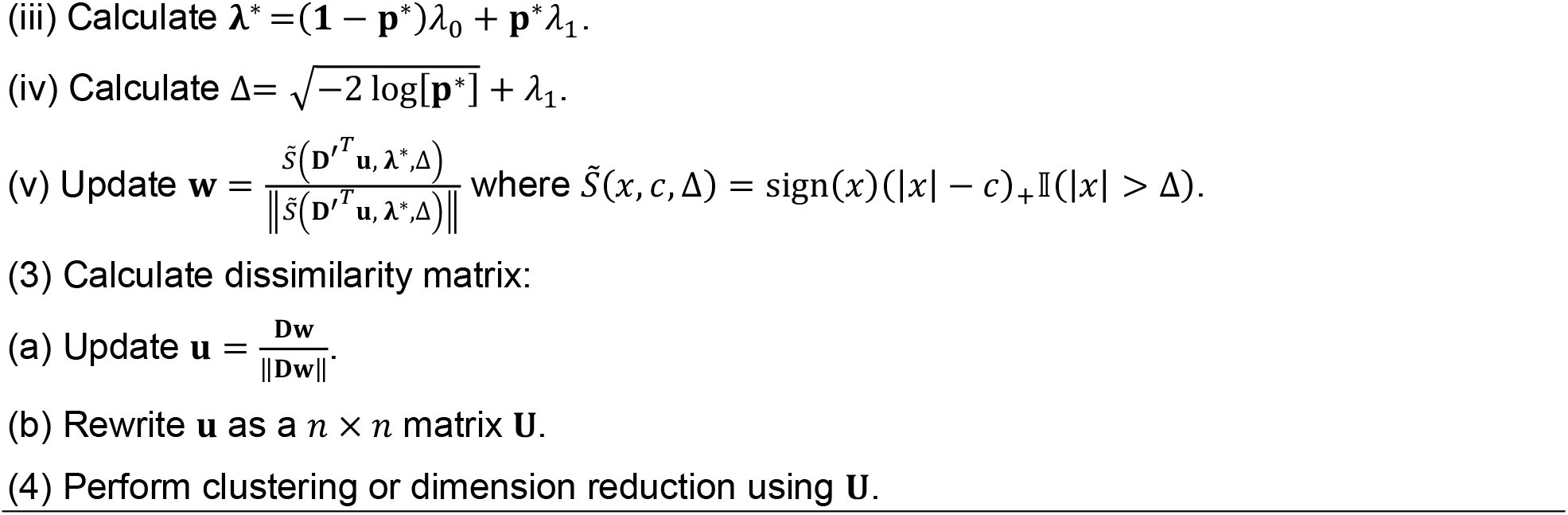

### Sparse hierarchical clustering with fused lasso

Here, we introduce the SHC-FL algorithm. An FL penalty contains L1 norms of both features and their successive differences^54^, which is defined as

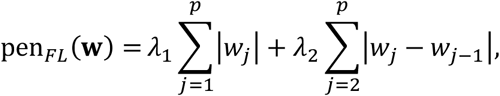

where λ_1_, λ^2^ ≥ 0 are hyperparameters. While λ_1_ encourages sparsity of features, λ^2^ encourages neighboring features to be similar and some to be identical.

We replace the lasso with the FL penalty in the objective of SHC, yielding the optimization criterion as

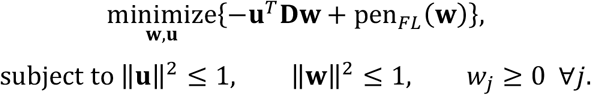

We iteratively optimize **u** and **w**. With **w** fixed, the optimal **u** is calculated in the same way as the one in SHC. With **u** fixed, the SHC-FL objective takes the form

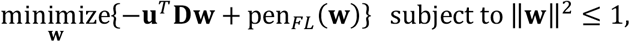

which is convex, but not smooth and not separable. To find stationary points of the above objective, we rewrite it as a Lagrangian function

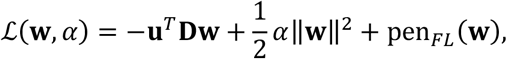

where α ≥ 0 is the Lagrange multiplier. The derivative of *L*(**w**, α) can be expressed as

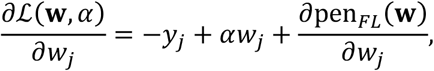

where **y** = **D**^*T*^**u**. At stationary points, the optimal *wj*’s needs to satisfy

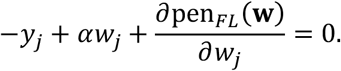

Let’s present the derivative of the FL penalty. Following the sub-differential calculus, the derivative of pen*FL* (**w**) is in the form

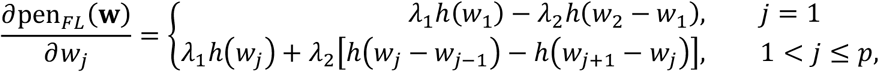

where *h*(*x*) is a subgradient of |*x*|, meaning *h*(*x*) = sign(*x*) when *x* ≠ 0 and *h*(*x*) ∈ [−_1, 1_]when *x* = 0. It is easy to see ∂pen_*FL*_ (α**w**)/∂(α*w*_*j*_) = ∂pen_*FL*_ (**w**)/∂*w*_*j*_, where α is a positive constant. Thus, a scaling of **w** does not affect the derivative of pen_*FL*_ (**w**).

Now, we solve for the optimal **w**. Let **w**′ = α**w**. Because 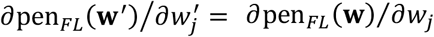, the necessary condition of the stationary points of **w** becomes

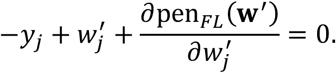

To solve **w**′ in the above equation, we construct an axillary convex optimization problem whose solution coincides with **w**′:

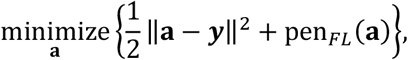

where **a** ∈ ℝ^*p*^. This is a signal approximation problem under the fused lasso penalty, with the solution called the fused lasso signal approximator (FLSA). We employ an FLSA solver proposed by Liu et al.^55^, because of its high computational efficiency given a pair of λ_1_ and λ^2^, and strong scalability in high dimension^56^. Once **w**′ is obtained, we get **w** = **w**′/‖**w**′‖ such that ‖**w**‖^2^ = _1_.

#### SHC-FL algorithm

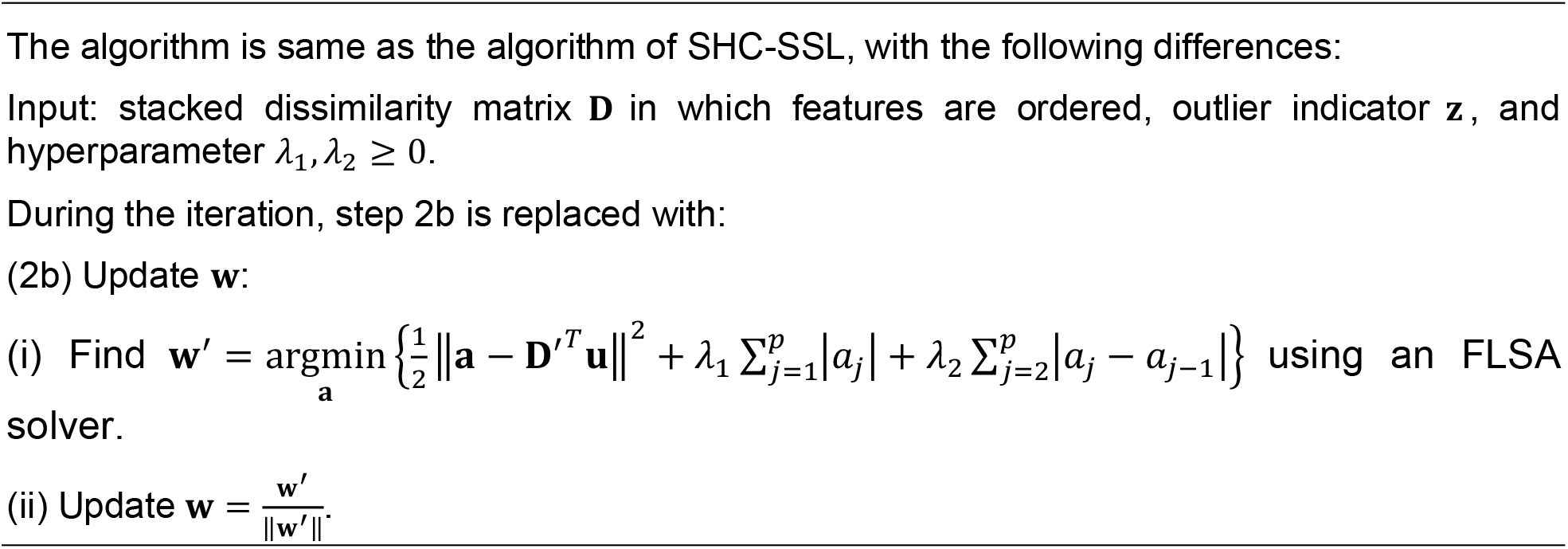

### Hyperparameter selection

We now turn to the selection of hyperparameters. In the above algorithms, the hyperparameters include: (1) *s* in SHC; (2) (λ_1_, λ^2^) in SHC-FL; and (3) λ_0_ in SHC-SSL, where we keep λ_1_ to a small constant. Let λ denote a single or a pair of hyperparameters in any one of the above algorithms. As in SHC, we employ a permutation-based approach, the gap statistic, to select the optimal value of λ^21,57^. Given a hyperparameter λ, the gap statistic measures the strength of the clustering based on real data with respect to the one based on randomly permuted data that are supposed to have no cluster. We quantify the strength of the clustering as

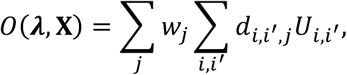

where **X** denotes the dataset used. Let **X** be the original real dataset, and **X**_1_, …, **X**_*B*_ denote *B* randomly permuted datasets by permuting samples within each feature. The gap statistic is calculated by

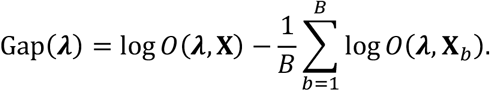

The optimal value of **λ** is obtained at the largest gap statistic.

In addition, to accelerate convergence of the presented algorithms for a series of *N* hyperparameter candidates {**λ**_1_, …, **λ**_*N*_}, we employ a “warm start” strategy: since solutions for similar hyperparameters are close, **w** is initialized as a solution for a nearby hyperparameter so that its solution can be found relatively quickly.

### Simulation data

We generated simulation datasets by adapting a simulation procedure described by Brodinová et al.^58^. Each dataset has *n* samples characterized by *p* features, with *p* = *p*_*d* +_ *p*_*n*_, where *p*_*d*_ is the number of discriminant features and *p*_*n*_ is the number of uninformative features. Each sample comes from one of *C* clusters. Let *n*_*t*_ be the size of the cluster *t*, and we have 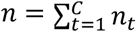. In this study, we choose *p*_*d*_ = 50, *p*_*n*_ = 950, *C* = 3, and *n*_*t*_ = 40 for *t* =1, 2, 3.

Samples of each cluster are fully characterized by the discriminant features, which follow a multivariate normal distribution with a mean vector 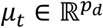 and a covariance matrix 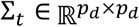, for *t* = _1_, …, *C*. The elements of the mean vector μ_*t*_ are constructed as

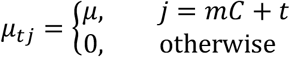

where 1 ≤ *j* ≤ *p*_*d*_, μ is a constant, and *m* is any nonnegative integer. The larger μ, the farther away the clusters are located geometrically. The covariance matrix Σ_*t*_ is constructed as

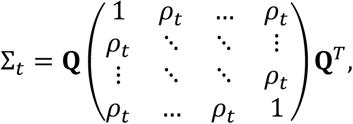

where **Q** is a random orthogonal matrix and ρ_*t*_ is a random number sampled from a uniform distribution *U*(*a*_ρ_, *b*_ρ_) with 0 < *a*_ρ_, *b*_ρ_ < 1. Here, we choose μ = 2, *a*_ρ_ = 0.1, and *b*_ρ_ = 0.9 to generate clusters that are close to each other, rendering a nontrivial problem for sparse clustering algorithms. For the uninformative features of each sample, we generate *p*_*n*_ random numbers that are independently and identically distributed from the standard normal distribution.

### Benchmarking of sparse hierarchical clustering algorithms using simulation data

Using the above procedure, we randomly generated 100 simulation datasets for benchmarking performances of the presented algorithms. In all the analyses, the dissimilarity metric between features was the squared distance, and the hyperparameters were chosen based on gap statistic profiles. In the SHC-SSL analyses, we set λ_1_ = 0.00_1_ and chose the optimal λ_0_ based on gap statistic profiles.

To evaluate the performances of the presented algorithms, we used the following performance metrics.

#### Recovery of discriminant features

To evaluate feature selection, given **w** from the result of the algorithms, we regard features {*j*|*w*_*j*_ > 0} as predicted discriminant features, and all the rest of features are predicted uninformative features. By comparing them with the true discriminant and uninformative features, we obtain precision, recall, and F1 score of the discriminant features.

#### Silhouette

To measure the clustering effect, we calculate the silhouette value using the dissimilarity matrix **U** and true cluster labels of samples. We use the silhouette value to measure the consistency of **U** from the true clusters rather than the consistency of a predicted partition from a fixed dissimilarity matrix.

#### Concordance of clusters along dendrogram after hierarchical clustering

The resulting dissimilarity matrix **U** is subjected to hierarchical clustering. To evaluate consistency of the resulting hierarchy of samples, represented as a dendrogram, with their true cluster labels, we calculate the concordance of cluster along the dendrogram as

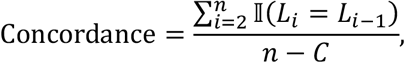

where *L*_*i*_ is the label of the cluster to which the sample *i* belongs. The denominator *n* − *C* normalizes the range of the concordance to be within 0 and 1. The larger the concordance, the better the clustering. In the case of the worst clustering, when successive samples always belong to different clusters, the concordance becomes 0; in the case of an ideal clustering, when all within-cluster samples are tightly connected, the concordance is 1.

This metric measures a combined effect of hierarchical clustering and the chosen linkage criterion. Given **U**, different linkage criteria generally produce different hierarchies. To compare different methods, we used the average linkage throughout this study.

#### Adjusted Rand index and purity of partitioned clusters after Leiden clustering

The resulting dissimilarity matrix **U** is used to find a shared *k* -nearest neighbor graph, where *k* = 20. The neighbor graph is then subjected to Leiden clustering, in which the resolution parameter is set as 1, to partition samples into clusters. To evaluate consistency of the partitioned clusters with the true cluster labels, we calculate adjusted Rand index and purity. The larger these metrics, the better the clustering.

### Pseudo-cell strategy for reducing computational cost of SHC-SSL

All SHC-based algorithms involve a dissimilarity matrix 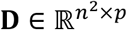, making the requirements of memory and computation scale quadratically with the sample size *n*. Thus, it is expensive to naïvely apply SHC-SSL when *n* is large. To reduce the computational cost, one can group cells with similar transcriptomes to form pseudo-cells, which are then subjected to SHC-SSL to select discriminant features. The selected features are used in the final clustering of all cells. An additional advantage of the pseudo-cell strategy is that it reduces technical noise and stabilizes variance, which may lead to more robust feature selection.

To construct pseudo-cells, specifically, one can use any efficient clustering algorithm, such as the Leiden algorithm, to group cells into many small clusters. Then, within each small cluster, pseudo-cells can be formed by calculating the mean of expressions of its constituent cells. This strategy is similar to the one used in Vision^59^, where pseudo-cells were termed “micro-clusters”. We employed this strategy for datasets with a large number of cells to reduce computational cost and select discriminant markers that best discriminate pseudo-cells.

### Neighborhood recurrence test

To test if a marker of a cell is recurrently up- or down-regulated in a biological subgroup rather than randomly perturbed, we devised a statistical test based on the idea that a recurrent marker should be enriched in the local neighborhood where the cell is located. The neighborhood recurrence test consists of two steps. First, we construct a K nearest neighbor graph of cells using their similarities based on discriminant markers. We define *S*_*k*_(*i*) to be the local neighborhood of cell *i*, for _1_ ≤ *i* ≤ *n*, where *S*_*k*_(*i*) includes the *k*-nearest neighbors of *i*. Second, we perform a statistical test to determine if marker *j* is significantly up- or down-regulated in the local neighborhood *S*_*k*_(*i*). Let μ_*j*_ and 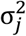 denote the mean and variance of the expression of marker *j*, which can be estimated as μ = ∑_*i*_ *X*_*ij*_ /*n* and 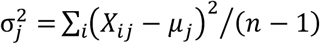, respectively. The mean expression of marker *j* for cells in the local neighborhood *S*_*k*_(*i*) can estimated as

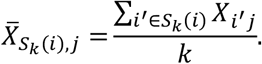

By the central limit theorem, when *k* is sufficiently large, 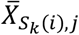 will be approximately normal distributed with its mean as μ_*j*_ and variance as 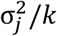. In our studies, we chose *k* = 20. Next, we calculate the neighborhood Z score of marker *j* for cell *i* as

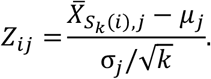

Thus, the distribution of the neighborhood Z score is approximately the standard normal distribution, i.e., *Pr*(*Z*) = *N*(0,1). Accordingly, the p-value of marker *j* that is recurrently up- or down-regulated in the local neighborhood of a cell *i* is *Pr*(*Z* > *Z*_*ij*_) or *Pr*(*Z* < *Z*_*ij*_), respectively. Finally, we adjust the p-values of all tested markers for each cell using the Benjamini-Hochberg procedure.

### Benchmark of hierarchical clustering performance of RECOMBINE markers and DEGs using biological datasets

We assessed the hierarchical clustering strength of RECOMBINE markers and DEGs across five biological datasets. For each dataset, UMAP projections were generated to visualize cell embeddings based on all genes, RECOMBINE markers, and DEGs. To quantify the hierarchical clustering strength of the RECOMBINE markers and DEGs, the gap statistic was computed in comparison to randomly selected gene sets of the same size,

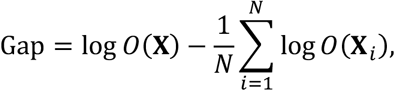

where **X** represent the expression matrix of the selected markers, **X**_*i*_ denote the expression matrix of a random gene set of the same size as **X**, and the objective function **X** is defined according to the sparse hierarchical clustering framework described earlier. For DEGs, gene weights were uniformly assigned and normalized to ensure an L2 norm of 1. The processing and analysis of each dataset are described below.

Data collection. For murine hematopoiesis, Dahlin et al., Blood (2018)^66^, the raw count matrices for six wild-type samples from the Lin^−^ Sca1^+^ Kit^+^ (LSK) and Lin^−^ Kit^+^ (LK) sorting gates were obtained from the dataset available at https://gottgens-lab.stemcells.cam.ac.uk/adultHSPC10X/. For murine hematopoiesis, Nestorowa et al., Blood (2016)^67^, log2 transformed counts and cell population annotations were downloaded from https://blood.stemcells.cam.ac.uk/single_cell_atlas. For murine hematopoiesis, Paul et al., Cell (2015)^68^, the raw counts data, and annotations of paul15_clusters were download through the Scanpy function sc.datasets.paul15(). For C. elegans embryogenesis, Packer et al., Science (2019)^69^, the expression data and development annotations of embryos were downloaded from https://depts.washington.edu/trapnell-lab/software/monocle3/celegans/data/.

#### Data preprocessing and analysis

For each dataset, low-quality cells were filtered out following the preprocessing steps described in the original publications, and the remaining data were log-normalized. PCA was performed with 20 PCs used in the downstream analysis. This was followed by batch correction across samples or batches using Harmony for the datasets of Dahlin et al. and Packer et al. Cell type annotations were obtained from the original publications if accessible, except that for the dataset of Dahlin et al cell type labels were assigned as 19 clusters identified by unsupervised clustering based on all genes. To generate pseudo-cells as mini-clusters, the Seurat function FindClusters was applied with a resolution of 50, and pseudo-cell expression profiles were computed as the average expression of cells within each mini-cluster. Discriminant markers were selected using RECOMBINE applied to the pseudo-cell expression profiles. DEGs were identified using the Seurat function FindAllMarkers, comparing clusters identified by FindClusters at a resolution of 2. A final list of DEGs was aggregated by selecting the top significant positive markers from each cluster, matching the total number of markers identified by RECOMBINE.

### Applying RECOMBINE to biological datasets for in-depth case studies

We applied RECOMBINE to bulk and single-cell omics datasets. In each analysis, we set λ_1_ = 0.000_1_, and chose the optimal λ_0_ based on gap statistic profile. The squared distance was used as the dissimilarity metric in SHC-SSL. The dissimilarity matrix based on discriminant markers was subjected to building a K nearest neighbor graph, where K = ^2^0, followed by Leiden clustering, where the choice of resolution parameter depends on the datasets. The cells were visualized in heatmaps showing Leiden clusters, where cells within each Leiden cluster were hierarchically clustered using the average linkage.

#### Selecting targeted panel from scRNA data for spatial molecular profiling of mouse visual cortex

For scRNA data, we downloaded an scRNA-seq dataset of mouse visual cortex from Allen Brain Atlas^26^ (http://celltypes.brain-map.org/api/v2/well_known_file_download/694413985) and filtered out cells with low quality. In total, we obtained 14,249 cells of 23 annotated cell types belonging to glutamatergic neurons, GABAergic neurons, and non-neuronal cells. We employed Seurat to normalize the data using the “LogNormalize” method and selected 10,000 highly variable genes using the “vst” method. Before applying RECOMBINE to the scRNA data, to reduce computational cost of SHC-SSL imposed by the large numbers of cells, we generated 287 pseudo-cells corresponding to clusters identified by Leiden clustering with a resolution of 50. Then, we used RECOMINE to select discriminant markers from 10,000 highly variable genes of these pseudo - cells. As a result, 366 genes were selected as discriminant markers of the mouse visual cortex scRNA data.

For spatial data, we downloaded a STARmap dataset of 1020 genes mapped in mouse primary visual cortex^12^ (https://www.dropbox.com/sh/f7ebheru1lbz91s/AADm6D54GSEFXB1feRy6OSASa/visual_1020/20180505_BY3_1kgenes). The spatial location coordinate of each cell was extracted from “labels.npz” according to the method provided by the original paper describing STARmap^12^. In total, there were 1,549 cells. We employed Seurat to normalize the data using the “LogNormalize” method with a scale factor of 100. After clustering, we annotated cell clusters into cell types based on expression of marker genes. Specifically, Slc17a7 and Gad1 were used to annotate excitatory and inhibitory neurons, respectively; Nov, Rorb, Sulf2, and Pcp4 were used to annotate eL2/3, eL4, eL5, and eL6 neurons, respectively; Vip, Sst, and Pvalb were used to annotate VIP, SST, and PVALB neurons, respectively; and Pdgfra, Aqp4, Enpp2, and Bsg were used to annotate microglia, astrocyte, oligodendrocyte, and endothelial cells, respectively. As the paired scRNA data of mouse visual cortex did not include microglia cells, we filtered out microglia cells from the spatial data, resulting in 1,486 spatially resolved cells remaining in our analysis. Of 366 discriminant markers identified from the scRNA data, 248 genes were mapped in the spatial data of mouse visual cortex. Next, we performed Leiden clustering of 1,486 cells from the spatial data using the 248 discriminant markers, where a resolution of 0.5 was used. Cells were annotated based on their markers and mapped to their physical locations. Finally, we performed RECOMBINE neighborhood recurrence tests to identify markers that characterize cell subpopulations in the spatial data.

#### Identifying markers that characterize cell states of CD8 tumor-infiltrating T cells across cancer types

We downloaded SingleCellExperiment object of scRNA-seq dataset of pan-cancer CD8 T cells (https://zenodo.org/record/5461803), which were obtained by integrating 234,550 tumor-infiltrating CD8 T cells of 316 patients across 21 cancer types from multiple sources^31^. To reduce noise, cells within each source had been partitioned into small groups (“miniclusters”). In total, the expression matrix had a dimension of 11,972 miniclusters by 11,772 genes. The downloaded data had been log_2_-transformed, and 1,500 highly variable genes were provided. Before applying RECOMBINE to the dataset, we generated pseudo-cells using Seurat to reduce computational cost of SHC-SSL. Specifically, the 1,500 highly variable genes were scaled and subjected to principal component analysis, from which the top 50 principal components were used for embedding. The embeddings were corrected using Harmony to mitigate batch effects across data sources. Then, shared nearest neighbor graphs were constructed, and Leiden clustering with a resolution of 30 was used to generate 205 pseudo-cells. We applied RECOMBINE to the pseudo-cell data to select discriminant markers and extract markers from the discriminant markers for all cells. In Leiden clustering of all cells based on the selected discriminant markers, the resolution parameter was set to 2, which identified 15 clusters, including naïve cells expressing TCF7 (c1); memory cells expressing IL7R, MALAT1, or FOXP4 (c2-5); resident memory cells expressing LGALS3 (c6); effector memory cells expressing GZMK, GZMA, or PFN1, (c7-9); exhausted cells simultaneously expressing high levels of PDCD1, CTLA4, TIGIT, and TOX (c10-11); interferon-stimulated genes (ISG)-positive cells (c12); natural killer (NK)-like cells expressing both NKG7 and KLRD1 (c13-14); and mucosal-associated invariant T cells (MAIT) expressing both KLRB1 and TMIGD2 (c15).

To study compositions of cell clusters associated with anti–PD-1 therapy response, we downloaded the clinical response data from the report by Sade-Feldman et al.^38^ (Supplementary Table 1, sheet patient-scRNA data) and used data of patients treated with anti–PD-1 therapy, where responders (N=7) included patients with complete and partial responses, and non-responders (N=14) included those with stable or progressive disease based on RECIST. The pan-cancer dataset included 3,853 CD8 T cells from these 21 patients. The statistical comparison of cell compositions between responders and non-responders was based on Wilcoxon rank sum test.

To validate the association between the composition of GZMK^+^HAVCR2^-^ EM T cells and anti-PD-1 response, we used a dataset of CD8 T cells from non-small-cell lung cancer patients (N=34) treated with anti–PD-1 therapy in the report by Liu et al.^39^, which was not included in the pan-cancer CD8 T cell dataset. We downloaded scRNA-seq, scTCR-seq, and metadata from the Gene Expression Omnibus (GEO; GSE179994), and response data from Liu et al.^39^ (Supplementary Table 1). Of 34 patients, 11 patients had post-treatment samples (N=14) with annotated clinical responses (9 responders vs. 5 non-responders). Based on the scTCR-seq and metadata, we extracted all CD8 T cells having TCR clonotypes shared by exhausted CD8 T cells, resulting in 13,403 cells defined as tumor-reactive CD8 T cells. After constructing 286 pseudo-cells, we applied RECOMBINE to this dataset and selected 236 discriminant genes. Leiden clustering of all tumor-reactive CD8 T cells with the discriminant genes identified 3 clusters: GZMK^+^HAVCR2^-^ EM, exhausted, and proliferative T cells. Finally, we compared compositions of GZMK^+^HAVCR2^-^ EM T cells from post-treatment samples between responders and non-responders after anti–PD-1 therapy.

#### Identifying markers that characterize a rare subpopulation of mouse intestine

We downloaded an scRNA-seq dataset of intestinal organoid cells from the GEO (GSE62270). The data were preprocessed as described previously^40^. Mitochondrial genes, ERCC spike-ins, and genes associated with clustering artifacts (Rn45s, Malat1, Kcnq1ot1, A630089N07Rik, and Gm17821) were excluded as described previously^40,62^. The resulting dataset had an expression matrix of 238 cells by 3,773 genes. Finally, log_2_ transformation was applied. Using this preprocessed dataset, we ran RECOMBINE to identify markers that discriminate both common and rare cell subpopulations. In Leiden clustering, the resolution parameter was set to 0.6.

#### Identifying markers that discriminate heterogeneous cells of human tissues in Tabula Sapiens

We downloaded h5ad files containing raw counts and cell ontology class annotations from the resource website (https://tabula-sapiens.sf.czbiohub.org/) of the Tabula Sapiens^70^. Data preprocessing and RECOMBINE were applied to each tissue individually before integrating the results for combined analysis. For each tissue, cells with fewer than 200 expressed genes, cells in the top 10th percentile of expressed gene counts, and cells with over 15% of transcripts originating from mitochondrial genes were excluded. The data were then log-normalized. Principal component analysis (PCA) was performed, selecting 30 principal components based on elbow plots, and batch effects across donors were corrected using Harmony. Mini-clusters were identified via unsupervised clustering using the Seurat FindClusters function with a resolution of 30, and pseudo-cell expression profiles were calculated as the average expression of cells within each mini-cluster. RECOMBINE was applied to the pseudo-cell expression data to select discriminant markers, followed by neighborhood recurrence tests to compute neighborhood Z-scores and adjusted p-values. RECOMBINE markers were extracted for each cell ontology class by selecting the top 10 markers with more than 50% of cells showing significant neighborhood recurrence and an average neighborhood Z-score greater than 2. Subsequently, the expression profiles of cells within the same ontology class were averaged to construct a pseudobulk expression matrix of cell ontology classes for each tissue. These pseudobulk matrices were pooled to generate an integrated expression matrix of cell ontology classes across all tissues based on RECOMBINE markers. Finally, UMAP projection and hierarchical clustering were performed to visualize the relationships among cell ontology classes and to uncover patterns of RECOMBINE markers across diverse cell types.

### Functional enrichment of discriminant markers and signature score calculation

To relate RECOMBINE-selected discriminant markers with biological functions, we performed functional enrichment using Enrichr^63^ with the MSigDB hallmark signatures^64^ and WikiPathways^65^ provided by the Enrichr web server. Let’s denote *e*_*ij*_ as the normalized and log-transformed expression of gene *j* in cell *i*, and *G* as the gene set of a signature. Following the signature score defined in Vision^59^, the signature score can be calculated as

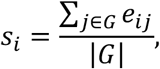

where |*G*| denotes the size of the gene set *G*. Since discriminant markers can optimally distinguish cells, we calculate the signature score using only discriminant markers as

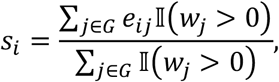

where II(*w*_*j*_ > 0) is an indicator function of gene *j* selected in discriminant markers. As reported by DeTomaso et al.^59^, even if the expression values are normalized, the signature scores calculated as described above may still depend on technical variability across cells. To remove this variability, the signature score can be normalized using the expected mean and variance of the score of a random gene set having the same number of genes. For a random gene set, the expected mean of the signature score in cell *i* is

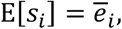

where 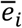 is the mean of expressions in cell *i*, and the expected variance of the signature score is

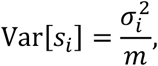

where 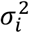 is the variance of the expressions in cell *i*, and *m* = ∑_*j*∈*G*_ II(*w*_*j*_ > 0). Finally, the signature score is normalized as

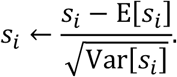

## Data availability

The simulation data are available and can be reproduced using the RECOMBINE R package. All the biological datasets are public and were found in previous papers as described in the Methods.

## Code availability

The RECOMBINE R package is available at https://github.com/korkutlab/recombine.

## Acknowledgements

We thank Lele Zhu for his comments on the pan-cancer CD8 T cell analysis. Editorial support was provided by Bryan Tutt, Scientific Editor, Research Medical Library at MD Anderson Cancer Center. This work is supported with grants from MDACC Support Grant P30 CA016672 (the Bioinformatics Shared Resource) (AK), U01CA253472 (AK).

## Author contributions

X.L. and A.K designed the study, developed the algorithms, performed data analyses, wrote the manuscript, J.N. performed data analyses of the Tabula Sapiens. All authors read and approved the final manuscript.

## Additional information

**Supplementary information** is available for this paper.

## References

1. Osumi-Sutherland, D. et al. Cell type ontologies of the Human Cell Atlas. Nature Cell Biology 23, 1129–1135 (2021).

2. Zhang, Y., Gao, S., Xia, J. & Liu, F. Hematopoietic hierarchy – an updated roadmap. Trends Cell Biol 28, 976–986 (2018).

3. Becker, W. R. et al. Single-cell analyses define a continuum of cell state and composition changes in the malignant transformation of polyps to colorectal cancer. Nature Genetics 54, 985–995 (2022).

4. Lange, M. et al. CellRank for directed single-cell fate mapping. Nature Methods 19, 159–170 (2022).

5. Kong, W. et al. Capybara: A computational tool to measure cell identity and fate transitions. Cell Stem Cell 29, 635-649.e11 (2022).

6. Aldridge, S. & Teichmann, S. A. Single cell transcriptomics comes of age. Nature Communications 11, 1–4 (2020).

7. Liu, Z. & Zhang, Z. Mapping cell types across human tissues. Science 376, 695–696 (2022).

8. Elmentaite, R., Domínguez Conde, C., Yang, L. & Teichmann, S. A. Single-cell atlases: shared and tissue-specific cell types across human organs. Nature Reviews Genetics 23, 395–410 (2022).

9. Rao, A., Barkley, D., França, G. S. & Yanai, I. Exploring tissue architecture using spatial transcriptomics. Nature 596, 211–220 (2021).

10. Longo, S. K., Guo, M. G., Ji, A. L. & Khavari, P. A. Integrating single-cell and spatial transcriptomics to elucidate intercellular tissue dynamics. Nature Reviews Genetics 22, 627–644 (2021).

11. Moffitt, J. R. et al. Molecular, spatial, and functional single-cell profiling of the hypothalamic preoptic region. Science 362, (2018).

12. Wang, X. et al. Three-dimensional intact-tissue sequencing of single-cell transcriptional states. Science 361, (2018).

13. He, S. et al. High-plex imaging of RNA and proteins at subcellular resolution in fixed tissue by spatial molecular imaging. Nature Biotechnology 40, 1794–1806 (2022).

14. Gohil, S. H., Iorgulescu, J. B., Braun, D. A., Keskin, D. B. & Livak, K. J. Applying high-dimensional single-cell technologies to the analysis of cancer immunotherapy. Nature Reviews Clinical Oncology 18, 244–256 (2020).

15. Luecken, M. D. & Theis, F. J. Current best practices in single-cell RNA-seq analysis: a tutorial. Mol Syst Biol 15, e8746 (2019).

16. Hao, Y. et al. Integrated analysis of multimodal single-cell data. Cell 184, 3573-3587.e29 (2021).

17. Lähnemann, D. et al. Eleven grand challenges in single-cell data science. Genome Biology 21, 1–35 (2020).

18. Baran, Y. et al. MetaCell: Analysis of single-cell RNA-seq data using K-nn graph partitions. Genome Biol 20, 1–19 (2019).

19. Persad, S. et al. SEACells infers transcriptional and epigenomic cellular states from single-cell genomics data. Nature Biotechnology 1–12 (2023).

20. Dann, E., Henderson, N. C., Teichmann, S. A., Morgan, M. D. & Marioni, J. C. Differential abundance testing on single-cell data using k-nearest neighbor graphs. Nature Biotechnology 40, 245–253 (2021).

21. Witten, D. M. & Tibshirani, R. A framework for feature selection in clustering. J Am Stat Assoc 105, 713–726 (2010).

26. Tasic, B. et al. Shared and distinct transcriptomic cell types across neocortical areas. Nature 563, 72–78 (2018).

27. Waldman, A. D., Fritz, J. M. & Lenardo, M. J. A guide to cancer immunotherapy: from T cell basic science to clinical practice. Nature Reviews Immunology 20, 651–668 (2020).

28. van der Leun, A. M., Thommen, D. S. & Schumacher, T. N. CD8+ T cell states in human cancer: insights from single-cell analysis. Nature Reviews Cancer 20, 218–232 (2020).

29. Chow, A., Perica, K., Klebanoff, C. A. & Wolchok, J. D. Clinical implications of T cell exhaustion for cancer immunotherapy. Nature Reviews Clinical Oncology 19, 775–790 (2022).

30. Lowery, F. J. et al. Molecular signatures of antitumor neoantigen-reactive T cells from metastatic human cancers. Science 375, 877–884 (2022).

31. Zheng, L. et al. Pan-cancer single-cell landscape of tumor-infiltrating T cells. Science 374, (2021).

32. Good, C. R. et al. An NK-like CAR T cell transition in CAR T cell dysfunction. Cell 184, 6081-6100.e26 (2021).

33. Sen, D. R. et al. The epigenetic landscape of T cell exhaustion. Science 354, 1165–1169 (2016).

34. Pauken, K. E. et al. Epigenetic stability of exhausted T cells limits durability of reinvigoration by PD-1 blockade. Science 354, 1160–1165 (2016).

35. Im, S. J. et al. Defining CD8+ T cells that provide the proliferative burst after PD-1 therapy. Nature 537, 417–421 (2016).

36. Siddiqui, I. et al. Intratumoral Tcf1+ PD-1+ CD8+ T Cells with Stem-like Properties Promote Tumor Control in Response to Vaccination and Checkpoint Blockade Immunotherapy. Immunity 50, 195-211.e10 (2019).

37. Huang, Q. et al. The primordial differentiation of tumor-specific memory CD8+ T cells as bona fide responders to PD-1/PD-L1 blockade in draining lymph nodes. Cell 185, 4049-4066.e25 (2022).

38. Sade-Feldman, M. et al. Defining T cell states associated with response to checkpoint immunotherapy in melanoma. Cell 175, 998-1013.e20 (2018).

39. Liu, B. et al. Temporal single-cell tracing reveals clonal revival and expansion of precursor exhausted T cells during anti-PD-1 therapy in lung cancer. Nature Cancer 3, 108–121 (2021).

40. Grün, D. et al. Single-cell messenger RNA sequencing reveals rare intestinal cell types. Nature 525, 251–255 (2015).

46. Liang, S. et al. Single-cell manifold-preserving feature selection for detecting rare cell populations. Nat Comput Sci 1, 374–384 (2021).

47. Dumitrascu, B., Villar, S., Mixon, D. G. & Engelhardt, B. E. Optimal marker gene selection for cell type discrimination in single cell analyses. Nature Communications 12, 1–8 (2021).

48. Vargo, A. H. S. & Gilbert, A. C. A rank-based marker selection method for high throughput scRNA-seq data. BMC Bioinformatics 21, 1–51 (2020).

49. Rood, J. E., Maartens, A., Hupalowska, A., Teichmann, S. A., & Regev, A. Impact of the Human Cell Atlas on medicine. Nature Medicine 28(12), 2486–2496. (2022).

50. Li, X. et al. (2022). Precision combination therapies based on recurrent oncogenic coalterations. Cancer Discovery, 12(6), 1542–1559. (2022).

51. Ročková, V. & George, E. I. The Spike-and-Slab LASSO. J Am Stat Assoc 113, 431–444 (2018).

52. Ročková, V. Bayesian estimation of sparse signals with a continuous spike-and-slab prior1. Ann Stat 46, 401–437 (2018).

53. Hastie, T., Tibshirani, R. & Wainwright, M. Statistical learning with sparsity: The lasso and generalizations. Statistical Learning with Sparsity: The Lasso and Generalizations (CRC Press, 2015).

54. Tibshirani, R., Saunders, M., Rosset, S., Zhu, J. & Knight, K. Sparsity and smoothness via the fused lasso. J R Stat Soc Series B Stat Methodol 67, 91–108 (2005).

55. Liu, J., Yuan, L. & Ye, J. An efficient algorithm for a class of fused Lasso problems. in Proceedings of the ACM SIGKDD International Conference on Knowledge Discovery and Data Mining 323–332 (2010).

56. Xin, B., Kawahara, Y., Wang, Y. & Gao, W. Efficient generalized fused lasso and its application to the diagnosis of Alzheimer’s disease. in Association for the Advancement of Artificial Intelligence (2014).

57. Tibshirani, R., Walther, G. & Hastie, T. Estimating the number of clusters in a dataset via the gap statistic. J R Stat Soc Series B Stat Methodol 63, 411–423 (2001).

58. Brodinová, Š., Filzmoser, P., Ortner, T., Breiteneder, C. & Rohm, M. Robust and sparse k-means clustering for high-dimensional data. Adv Data Anal Classif 13, 905–932 (2019).

59. DeTomaso, D. et al. Functional interpretation of single cell similarity maps. Nat Commun 10, 1–11 (2019).

62. Gehart, H. et al. Identification of enteroendocrine regulators by real-time single-cell differentiation mapping. Cell 176, 1158-1173.e16 (2019).

63. Chen, E. Y. et al. Enrichr: Interactive and collaborative HTML5 gene list enrichment analysis tool. BMC Bioinformatics 14, (2013).

64. Liberzon, A. et al. The molecular signatures database hallmark gene set collection. Cell Syst 1, 417–425 (2015).

65. Kelder, T., et al. WikiPathways: building research communities on biological pathways. Nucleic Acids Research 40, D1301–D1307 (2012).

66. Dahlin, J. S., et al. A single-cell hematopoietic landscape resolves 8 lineage trajectories and defects in Kit mutant mice. Blood 131, e1–e11 (2018).

67. Nestorowa, S., et al. A single-cell resolution map of mouse hematopoietic stem and progenitor cell differentiation. Blood 128, e20–e31 (2016).

68. Paul, F., et al. Transcriptional heterogeneity and lineage commitment in myeloid progenitors. Cell 163, 1663–1677 (2015).

69. Packer, J. S., et al. A lineage-resolved molecular atlas of C. elegans embryogenesis at single-cell resolution. Science 365, eaax1971 (2019).

70. Tabula Sapiens Consortium, et al. The Tabula Sapiens: A multiple-organ, single-cell transcriptomic atlas of humans. Science 376, eabl4896 (2022).

