## Supplementary material for "Recurrent Composite Markers of Cell Types and States": Suppl. Figures

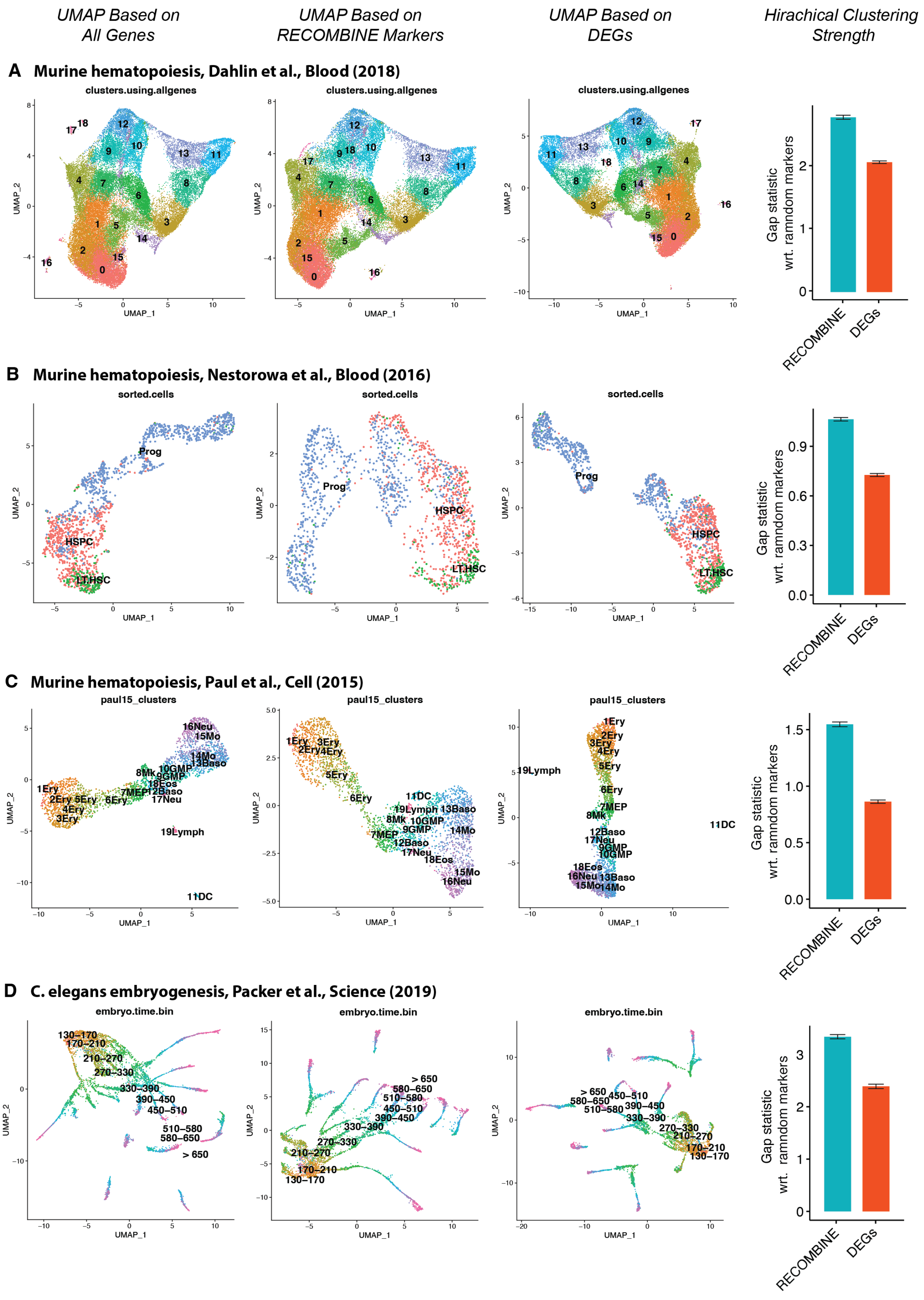
UMAP Based on All Genes
UMAP Based on RECOMBINE Markers
UMAP Based on DEGs
Hirachical Clustering Strength
A Murine hematopoiesis, Dahlin et al., Blood (2018)
clusters.using.allgenes
UMAP_1
UMAP_2
Gap statistic wrt. ramndom markers
RECOMBINE
DEGs
B Murine hematopoiesis, Nestorowa et al., Blood (2016)
sorted.cells
Prog
HSPC
LT.HSC
C Murine hematopoiesis, Paul et al., Cell (2015)
paul15_clusters
D C. elegans embryogenesis, Packer et al., Science (2019)
embryo.time.bin

**Figure S1. Benchmarking of hierarchical clustering performance of RECOMBINE markers compared with DEGs. (A-D)** 4 biological datasets. From left to right, UMAPs based on all genes, RECOMBINE markers, and top DEGs of the same size as RECOMBINE markers, are shown, and the last column evaluates hierarchical clustering performance by cell hierarchy discriminability (gap statistic) defined as the difference of hierarchical clustering strength between selected features versus random features of the same size.

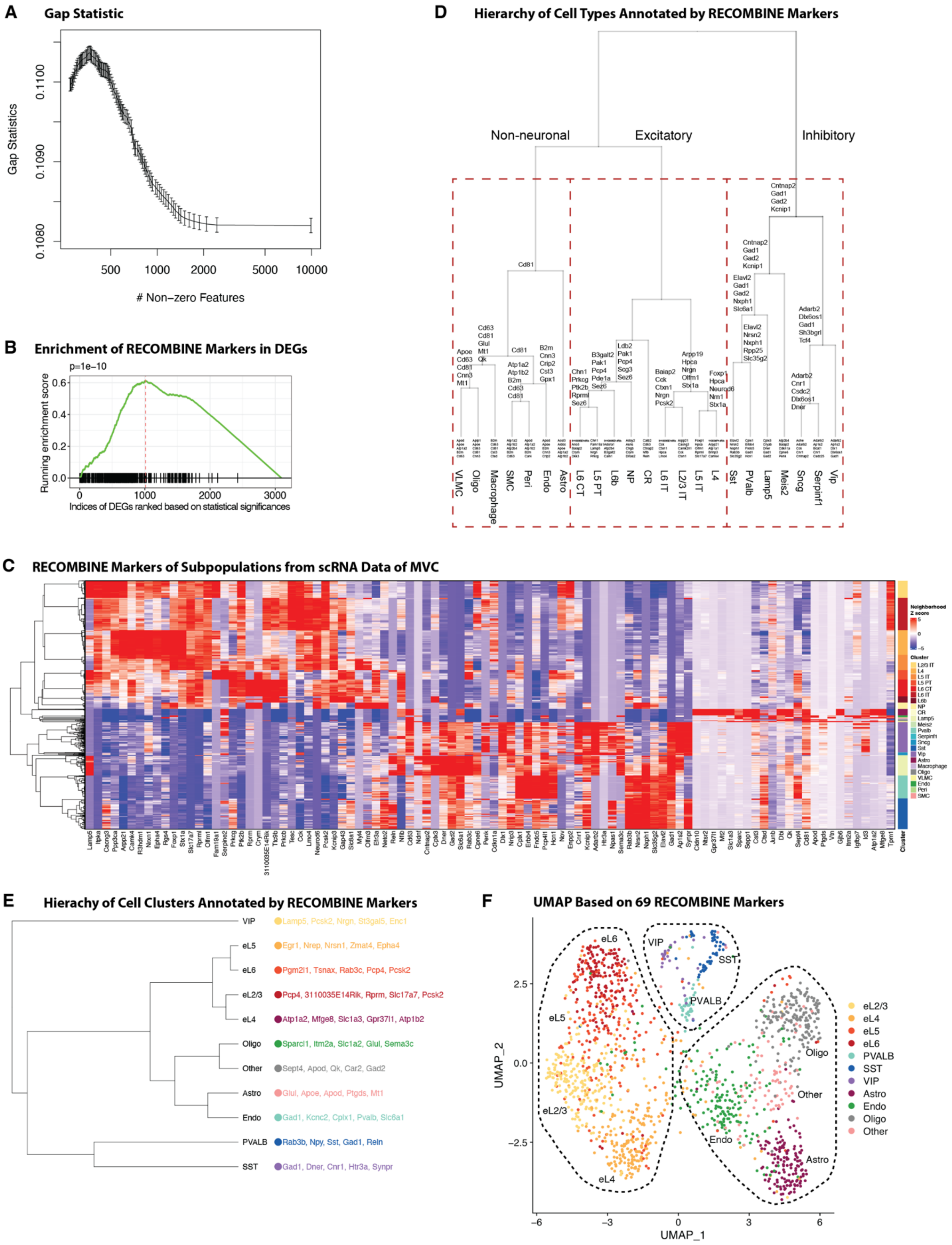
A
Gap Statistic
Gap Statistics
0.1080
0.1090
0.1100
500
1000
2000
5000
10000
### Non−zero Features
B
Enrichment of RECOMBINE Markers in DEGs
p=1e−10
Running enrichment score
0.0
0.2
0.4
0.6
0
1000
2000
3000
Indices of DEGs ranked based on statistical significances
D
Hierarchy of Cell Types Annotated by RECOMBINE Markers
Non-neuronal
Excitatory
Inhibitory
VLMC
Oligo
Macrophage
SMC
Peri
Endo
Astro
L6 CT
L5 PT
L6b
NP
CR
L6 IT
L2/3 IT
L5 IT
L4
Sst
PValb
Lamp5
Meis2
Sncg
Serpinf1
Vip
C
RECOMBINE Markers of Subpopulations from scRNA Data of MVC
Neighborhood Z score
Cluster
E
Hierachy of Cell Clusters Annotated by RECOMBINE Markers
VIP Lamp5, Pcsk2, Nrgn, St3gal5, Enc1
eL5 Egr1, Nrep, Nrsn1, Zmat4, Epha4
eL6 Pgm2l1, Tsnax, Rab3c, Pcp4, Pcsk2
eL2/3 Pcp4, 3110035E14Rik, Rprm, Slc17a7, Pcsk2
eL4 Atp1a2, Mfge8, Slc1a3, Gpr37l1, Atp1b2
Oligo Sparcl1, Itm2a, Slc1a2, Glul, Sema3c
Other Sept4, Apod, Qk, Car2, Gad2
Astro Glul, Apoe, Apod, Ptgds, Mt1
Endo Gad1, Kcnc2, Cplx1, Pvalb, Slc6a1
PVALB Rab3b, Npy, Sst, Gad1, Reln
SST Gad1, Dner, Cnr1, Htr3a, Synpr
F
UMAP Based on 69 RECOMBINE Markers
UMAP_2
UMAP_1
eL6
VIP
SST
PVALB
eL5
eL2/3
eL4
Oligo
Other
Endo
Astro
eL2/3
eL4
eL5
eL6
PVALB
SST
VIP
Astro
Endo
Oligo
Other

**Figure S2. RECOMBINE applied to scRNA-seq and STARmap data of mouse visual cortex.** **(A)** Gap statistic as a function of the number of selected genes. **(B)** Enrichment of RECOMBINE markers within the DEGs ranked by their significance levels (Wilcoxon test). **(C)** Heatmap of neighborhood Z scores showing gene modules across cell types of the scRNA data. **(D)** Hierarchical structure of cell types with each node annotated by RECOMBINE markers. **(E)** Hierarchy of cell clusters annotated by top 5 RECOMBINE markers. **(F)** UMAP of spatially resolved cells based on a reduced-size panel of RECOMBINE markers (N=69).

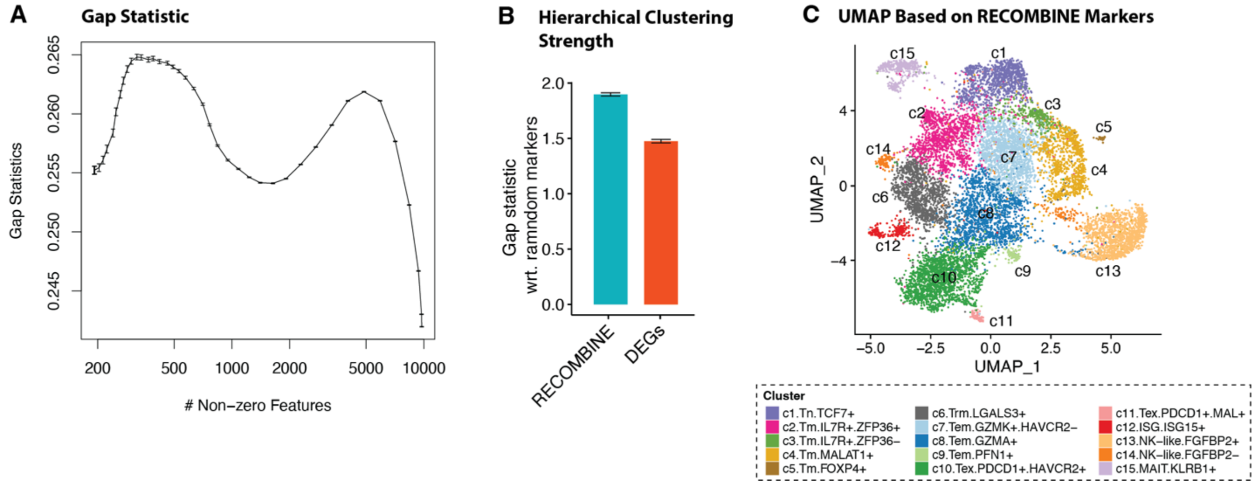


**Figure S3. RECOMBINE applied to scRNA-seq data of pan-cancer CD8 T cells. (A)** Gap statistic as a function of the number of selected genes. **(B)** Comparison of hierarchical clustering performance between RECOMBINE markers and top DEGs of the same size. **(C)** UMAP based on RECOMBINE markers and colored by clusters.

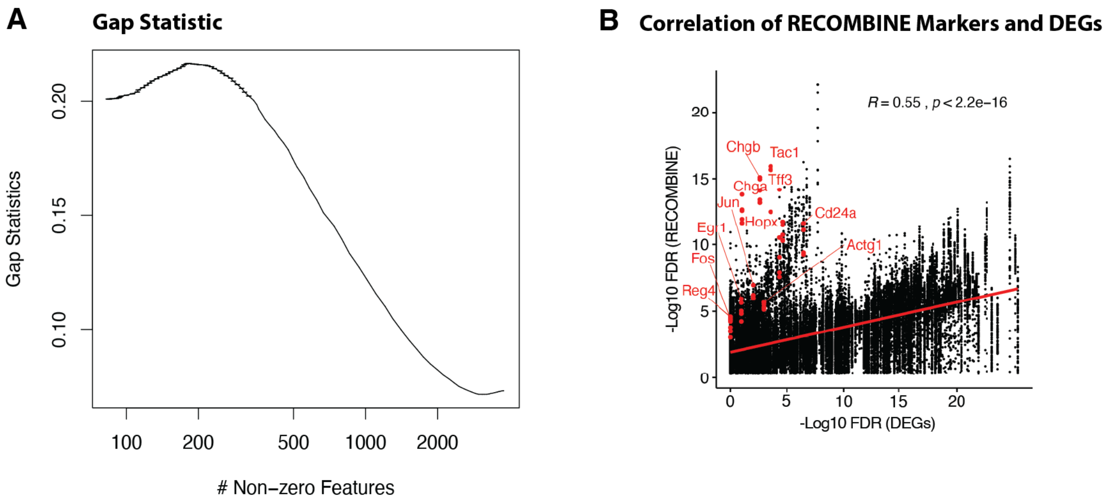


**Figure S4. RECOMBINE applied to scRNA-seq data of mouse intestine. (A)** Gap statistic as a function of the number of selected features. **(B)** Comparison of FDRs between RECOMBINE and DEGs for all cells. As FDRs of DEGs were obtained at the cluster level, we used the same FDRs for cells within the same cluster. The rare Reg4$^+$ subpopulation's markers of cluster c5 are highlighted in red.
